# A soluble host signal drives rapid, brain-predominant capsular thickening in *Streptococcus pneumoniae* via a putative sodium-dependent transporter (SPD_0642) and capsular prepromoter sequence

**DOI:** 10.64898/2026.03.28.714961

**Authors:** Nikola Tomov, Sabrina Hupp, Annelies Müller, Dario Baronti, Janine Lux, Irene Trillo, Cebile Lekhuleni, Tim van Opijnen, Federico Rosconi, Anne von Gottberg, Lucy J. Hathaway, Asparouh I. Iliev

## Abstract

The Streptococcus pneumoniae capsule is a major determinant of virulence, yet whether bacteria actively remodel it during infection remains unclear. Studying Swiss and South African clinical isolates (serotypes 1, 6B, 8, 12F, 19F, and 35B), we identified a rapid, tissue-specific response: capsule thickness increased within hours upon co-exposure to host cells and tissues. Only two 12F strains failed to thicken. Thickening was greatest in brain tissue, moderate in serum, and absent on the epithelium. This adaptation occurred independently of cod locus phase variation and nutritional factors, and was instead driven by a soluble, thermostable host signal (<3 kDa). Thickening correlated with neuroinflammation but did not require it, as it also occurred in contact with resting brain immune cells. It exacerbated meningitis in mice and enhanced bacteremia. Once induced, capsule thickening dampened inflammatory responses, coinciding with downregulation of pneumolysin, a major pro-inflammatory toxin. Genetic analysis of the non-thickening 12F isolates, together with targeted mutagenesis, identified two independent determinants of capsule-thickness modulation: a specific promoter-proximal element and SPD_0642, a conserved putative transporter encoded outside the capsule operon. Both contributed to the host-induced thickening phenotype. Pneumococci therefore rapidly remodel their surface in response to tissue-specific cues within the host, in a manner distinct from stochastic phase variation outside it.

**Importance:** Many bacteria are covered by a slimy outer layer, known as a capsule, that helps them evade the immune system. The amount of this layer can influence how easily harmful bacteria cause disease. Until now, scientists knew that bacteria can turn capsule production on or off through changes in their DNA. In this study, we show that Streptococcus pneumoniae, a common cause of serious infections, can also adjust its capsule in another way. It senses soluble signals from the tissues it enters, allowing it to recognize where it is in the body and to gradually change the thickness of its protective outer layer. This finding offers a new way of understanding how bacterial infections develop and may point to new treatment strategies.

## Introduction

*Streptococcus pneumoniae* is a globally prevalent pathogen that colonizes the human upper respiratory tract asymptomatically but can cause life-threatening pneumonia and other invasive pneumococcal diseases, such as meningitis and sepsis. Pneumococcal meningitis is a severe infection of the brain meninges associated with neurological complications such as cognitive deficits, learning disabilities or hearing loss (1). In 30% of all meningitis cases, the outcome is fatal due to severe complications such as brain swelling and ischemic damage(2). The most vulnerable sections of the population are infants (<5 years of age), elderly individuals, and immunocompromised patients with underlying diseases such as HIV (3). Mortality rates reach 50% in these groups (4, 5). Even vaccination cannot prevent such fatalities completely due to serotype shift(6).

A major virulence factor of *S. pneumoniae* is the polysaccharide capsule, which, based on its biochemical structure, defines the pneumococcal serotype. More than 100 different pneumococcal serotypes are known (7), some of which are associated with particularly severe disease (8, 9). Different hypotheses exist as to why one serotype causes more severe disease than the other. In a previous study, we described differences between serotypes in their ability to grow, not only in conventional lab media (10) but also in human cerebrospinal fluid (11), suggesting a competitive advantage for some serotypes. Another aspect is the difference in capsular thickness between serotypes, which may contribute to disease severity, as shown previously *in vivo* (12) — thicker capsules lead to a more severe course.

Bacteria (including *S. pneumoniae*) have developed capsule-modulating strategies to optimally colonize various host niches. Known as phase variation, it includes complex genetic reorganization, leading to switching capsular expression on and off (13). Phase variants differ in the capsular amount(14). Both variants coexist, and after being cultured on agar plates, two types of colonies are identified: opaque (higher capsule-producing properties, on-genotype) and transparent (lower capsule-producing properties, off-genotype) colonies. Opaques are better invaders and transparent better colonizers in animal models(15, 16), although all these analyses rely on preselected colonies on agar, which can eventually secondarily switch again. Currently, it is known that colony opacity is determined by the *cod* locus, which contains several components, such as the HsdM DNA methylase, which epigenetically controls the expression of capsule-synthesizing enzymes(14). Specifically, inversion of a fragment from a critical DNA-recognizing protein, HsdSa, that complements HsdM alters the target specificity of the HsdM methylase, thereby switching gene expression states(17). Other genetic regulators of bacterial capsule expression have also been identified, but their role in phase variation remains unknown.

Capsular genes are organized in an operon with a core promoter(18). Regulatory sequences such as the 37-CE fragment and others are positioned upstream of the core promoter and can also influence various aspects of the function of the operon(19).

Beyond capsule-mediated evasion, additional virulence determinants, such as cholesterol-dependent pore-forming cytolysin pneumolysin (PLY), further modulate host‒-pathogen interactions(20). At sublytic doses (normally present during disease), PLY can trigger cell stress responses, cytoskeletal rearrangement, and occasionally apoptosis depending on the cell type(21, 22). The released PLY also contributes to the development of the proinflammatory response(23).

In the brain, which is a unique immunological niche of the body with limited access to the acquired immune system, microglia, supported by astrocytes, represent the first line of defense. In the course of systemic bacterial infections in humans, multiple cytokines are released, leading to subsequent neutrophil infiltration. The proinflammatory factors TNF-α and IL-6 have major predictive value for the course of the disease(24). IL-8 (functional murine homologs of IL-8 are CXCL1 (KC) and CXCL2 (MIP-2)) is critically important for neutrophil recruitment from the blood to the brain and the accumulation of leukocytes at the site of infection(25).

In this study, we aimed to determine the dynamic changes in capsular thickness in various host niches, its correlation with the proinflammatory cytokine response and disease severity, and genetic factors determining its control in several recent clinical isolates from Swiss and South African patients.

## Results

### Epidemiological data of the strains

We selected six isolates from South African meningitis patients (serotypes 8, 12F, 19F and 35B) and two from Swiss airway infection patients (serotypes 1 and 6B). Serotype 1 is often used as a reference in epidemiological studies; 8 and 12F are the most common isolates in South Africa(26); 19F and 6B are associated with higher case fatality rates (27); and 35B is an emerging nonvaccine strain with an increased incidence of multidrug resistance (28). All strains (except the 35B, which is included only in the recent 21-valent pneumococcal conjugate vaccine) are included in the current 20-valent pneumococcal conjugate vaccine and 23-valent pneumococcal polysaccharide pneumococcal vaccines, whereas 8 and 12F are missing from the earlier 13- and 15-valent vaccines (29). Specific information about the patients from whom the strains were isolated is presented in Supplementary table T1.

### Capsular thickness increases after coincubation with glia, glia-conditioned medium, or FCS

We used a previously described FITC-dextran (2,000 kD) approach for bacterial capsule thickness measurement(10) (Fig. 1). Initially, we measured the total bacterial diameters, which correlated with capsular thickening (see Methods) on a widefield microscope (data in pixels), switching thereafter to a direct measurement on a confocal, subtracting the negative contrast image from the transmission image (Supplementary Fig. S1A, B). In these later experiments, we presented the results as absolute (µm) or as relative thicknesses (versus the mock-control) depending on the aim. Bacteria were initially cultured until OD_600_=0.6 in brain heart infusion broth (BHI). Their capsular thickness remained unchanged throughout all phases of multiplication (Fig. 1A). Coincubation with glia in serum-free medium significantly increased the capsular thickness in all strains except for the serotype 12F strain (50943) (Fig. 1B, Supplementary Fig. S2). To verify that the effect was indeed only capsule thickness specific, we measured the FITC-based negative contrast diameter, subtracting the size of bacteria (transmission image), confirming a significant increase proportional to the FITC-exclusion method (Supplementary Fig. S1A, B). Next, strain 50943 was serotyped, indicating the presence of a capsule. High-resolution negative contrast FITC-dextran imaging, compared with the capsule-deficient D39 strain (ΔCps (D39)), confirmed that the capsule was indeed present (Supplementary Fig. S1C). In the initial experiments, we used total bacterial size for thickening assessment; in later experiments, we used the exact measurement of bacterial thickness via confocal microscopy.

**Figure 1.**
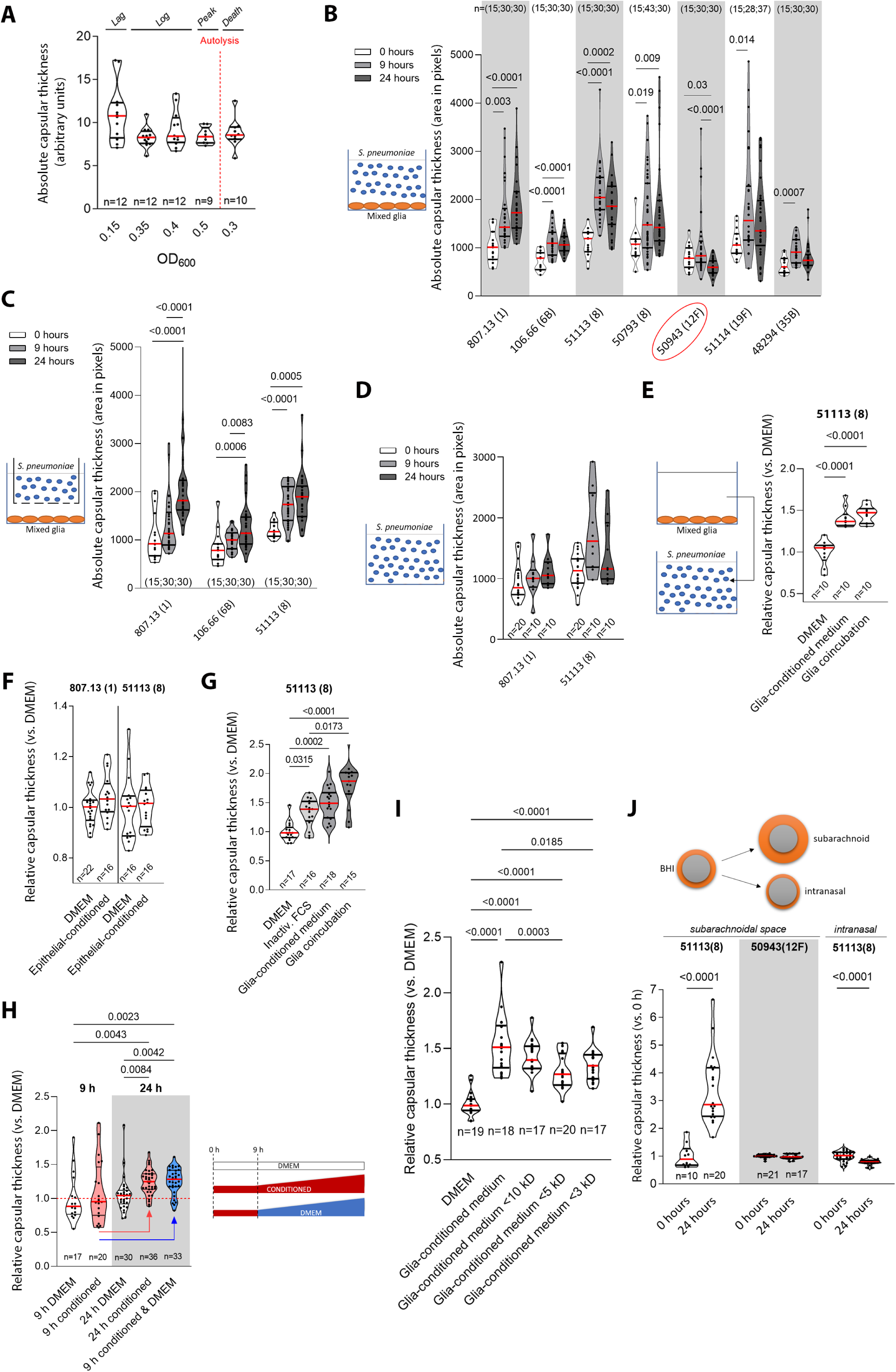
Host cell-mediated increase in capsule thickness. A. Capsular thickness in different phases of bacterial growth does not differ. B. Capsular thickness increases for all studied strains except 50943 (serotype 12F; circled in red) after 9 and 24 hours of direct coincubation with mixed glial cells. C. Host cell-mediated increase in capsule thickness at 9 and 24 h in indirect interaction between bacteria and glial cells through a semipermeable membrane. D. Unchanged capsular thickness at 9 and 24 h compared to 0 h for the 807.13 and 51113 strains in pure serum-free DMEM. E. Relative increase in capsular thickness (compared to nonconditioned DMEM) in the 51113 strain when preconditioned medium from nonactivated glial cells is used for 24 h. The effect is comparable to the increase in capsular thickness after direct coincubation with glia. F. Unchanged relative capsular thickness (compared to DMEM) of the 807.13 and 51113 strains after 24 h of incubation in epithelial cell (Detroit)-conditioned medium. G. With inactivated FCS, the capsular thickness increases at 24 h, but significantly less than that after direct glia coincubation. H. Relative increase in the capsular size of 51113 pneumococci incubated for 6 h and 24 h in pure DMEM, in glia-conditioned DMEM or for 6 h in conditioned medium, after which the mixture is exchanged for pure DMEM; the conditioned medium is sufficient for initiating a capsular increase at 24 h but not at 6 h. I. Capsular thickening capacity of glia-conditioned medium with a cutoff of higher molecular weight fractions, indicating that thickening is present even with a 3 kDa cutoff, indicating that the thickening factor is smaller than 3 kDa. Each symbol represents one image containing at least four bacteria. Each experiment is performed in duplicate three times on three different days, and the data are pooled (n=counted fields). J. When injected into the subarachnoid space after growth in BHI, the capsule-thickening 51113 strain exhibits substantial increases in capsular thickness. In contrast, the capsule of the in vitro nonthickening strain 50943 does not thicken during meningitis. Following intranasal colonization, however, the 51113 strain demonstrates a reduced capsular layer thickness. Each symbol represents one image containing at least four bacteria. Three animals per condition are analyzed, and an equal number of frames per animal per group (between 3 and 9) are collected and pooled together. Violin plots represent the median (red line) and quartiles (black line) (n=counted fields). All strains are compared via Mann-Whitney U-test or Kruskal–Wallis with Dunn’s multiple comparisons, and the p value is presented if significant.

The incubation of bacteria without direct contact with glial cells through a semipermeable membrane overnight induced a similar increase in capsular thickness (Fig. 1C). The incubation of bacteria with pure serum-free medium without glia did not change their capsule thickness (Fig. 1D). Incubation in medium conditioned beforehand with resting glia caused an increase in capsular thickness, which was identical to that observed with direct coincubation (Fig. 1E). Bacterial incubation in medium conditioned with epithelial cells failed to increase the capsular thickness (Fig. 1F). Incubation in inactivated fetal calf serum (FCS) thickened the capsule but not as much as that of live glial cells (Fig. 1G).

The increase in capsular thickness triggered by glia was due either to a signal from glial cells or to metabolic input. To test this hypothesis, we incubated strain 51113 with glia-conditioned medium for 6 h (Fig. 1H). No thickening during this period was observed before the medium was changed to nonconditioned DMEM. Twenty-four hours later, capsular thickening did occur, indicating that the factor in the conditioned medium is not metabolically required but acts as a necessary permissive signal early on (Fig. 1H). Furthermore, the results indicated that DMEM alone contains all the factors needed metabolically to support increased capsular thickness.

To study the density of the thickened capsule, we incubated thickened and nonthickened D39 pneumococci with a mixture of differentially fluorescently stained dextrans (70 and 2,000 kDa). In nonthickened bacteria, which both overlapped (homogenous capsule), in bacteria with thickened capsules following glial-conditioned medium, both dextrans demonstrated heterogeneous appearances, excluding the 2,000 kDa dextran but not the 70 kDa dextran (Supplementary Fig. S3).

### The thickening factor is smaller than 3 kDa, it is not a defensin, and it is heat stable

We fractionated glia-conditioned medium to exclude fragments above 10, 5 and 3 kDa via size exclusion filters (Fig. 1I). Capsular thickening was present even after the lowest 3 kDa cutoff size filter, indicating that the thickening factor was smaller in size. Some reduction in thickening was probably due to incomplete recovery of the factor in the ultrafiltrate.

Defensins are natural antibacterial factors with sizes between 3 and 5 kDa that are primarily secreted by neutrophils and epithelial cells and attack the bacterial cell wall/membrane. Nevertheless, some evidence indicates their release from glial cells. We exposed pneumococci to different concentrations of human β-defensin 3 (hBD3) at concentrations of 0.04, 0.6 and 5 µg/ml for 20 h without affecting pneumococcal capsular thickness (not shown). Heating at 95°C for 60 min did not alter the thickening capsular effect of the conditioned medium (not shown).

### Capsular thickness changes in animals corresponded to cell culture findings

To verify the relevance of the cell culture findings to systemic conditions, we injected the thickening 51113 strain and the nonthickening 50943 strain subarachnoid with 10^5 CFU to initiate meningitis in C57BL/6JRj mice (Fig. 1J). Twenty-four hours later, 51113 demonstrated capsular thickening, whereas 50943 remained unchanged. Intranasal colonization with the 51113 strain even resulted in reduced capsular thickness in the recovered nasal lavage fluid compared with the thickness of the bacterial stock in BHI used for the experiments (Fig. 1J). The lavage fluid contained an abundant number of *S. pneumoniae* colonies on blood agar, as an equivalent number of pneumococci was also identifiable under a microscope.

### When present, capsular thickening correlates with the neuroinflammatory response

We addressed the pathogenic importance of capsular thickening by studying the release of the proinflammatory cytokines TNF-α and IL-6 and the chemokine CXCL2 (Fig. 2A-C). Coculturing each strain in identical numbers (10^7 CFU/ml) with mixed glia elevated inflammatory cytokine release (Fig. 2A-C). The initial bacterial size (corresponding to capsular thickness; see Methods) correlated best with TNF-α release (Fig. 2D). The nonthickening strain 50943 (12F) remained an outlier in the chemokine correlation (Fig. 2E). Excluding the 12F strain, the correlations for all cytokines were significant at 0 h and 24 h (Fig. 2F). Thus, both initial capsular thickness and subsequent thickening by glia correlated well with the neuroinflammatory response in thickening strains.

**Figure 2.**
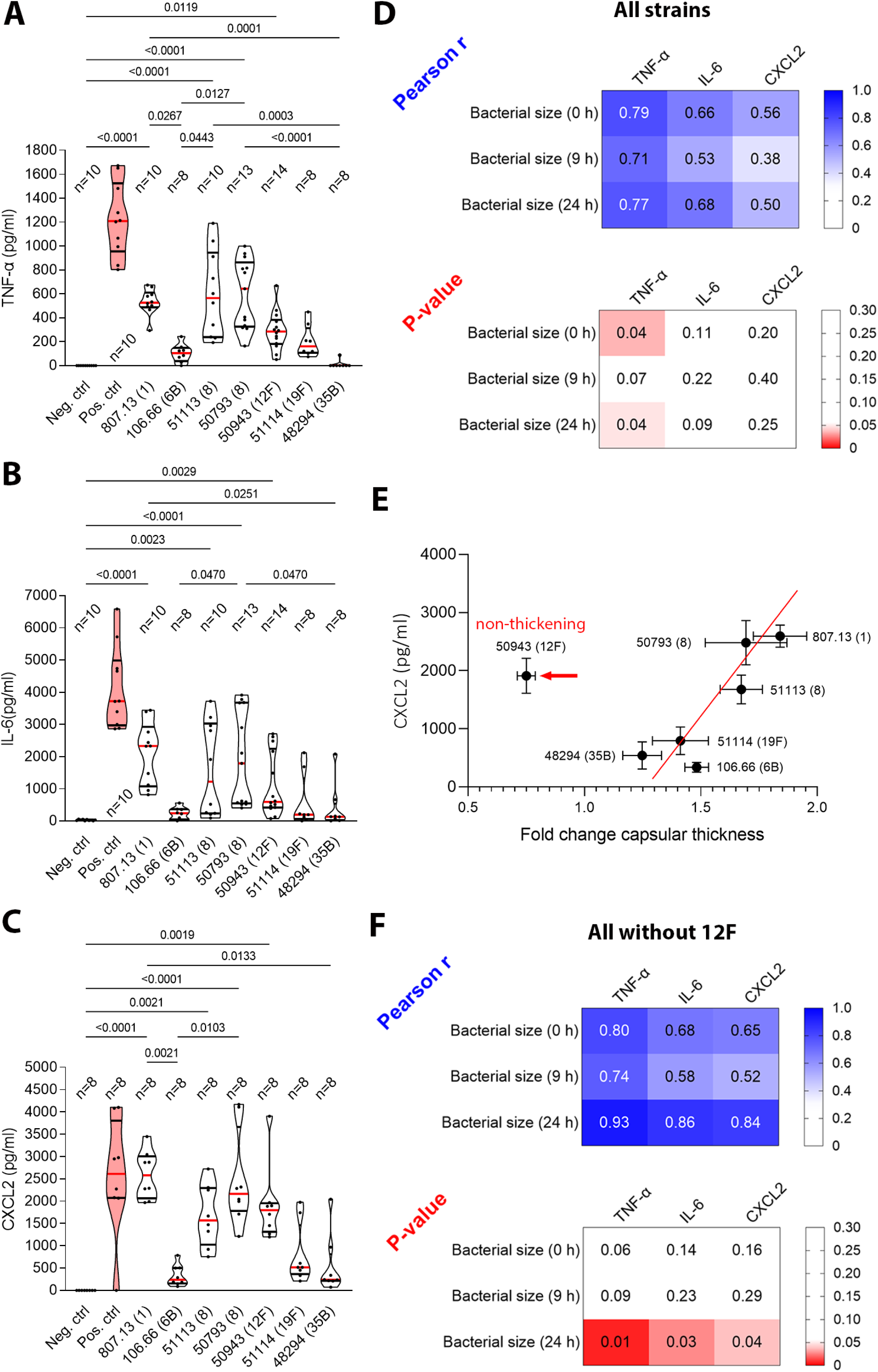
Cytokine release after strain coincubation with glia. A. TNF-α release (ELISA). B. IL-6 release (ELISA). C. CXCL2 release (ELISA) in mixed primary glial cultures after 24 h of direct coincubation with selected pneumococcal strains. No clear serotype- or strain-specific pattern is observed. All strains tested induce strong cytokine responses that are significantly greater than those of the negative (vehicle) control. A positive control (cells + 0.1 µg/ml lipopolysaccharide (LPS) (in red)) is included for all the assays. All violin plots represent the median (red) and quartiles (black). All strains are compared via Kruskal–Wallis with Dunn’s multiple comparisons, and the p value is presented if significant. Each point represents a biological replicate. D. Pearson’s R as an indicator of correlation (r>0.7) between bacterial size (FITC-dextran negative contrast as a capsular thickness correlate) and cytokine release for all strains tested. For all strains, a strong correlation is observed only with TNF-α. The p-value between capsular thickness and cytokine release for all strains tested is considered significant at p<0.05 (outlined in red). E. Linear correlation between changes in capsular thickness and CXCL2 release for all strains tested, except for the 12F strain. Data show means ± SEM. F. Pearson’s R as an indicator of correlation (r>0.7) between bacterial size (FITC-dextran negative contrast) and cytokine release for all strains tested when excluding the outlier 12F strain. A strong correlation is observed for all the tested cytokines. The P value between capsular thickness and cytokine release for all strains tested is considered significant at p<0.05 (outlined in red).

### Capsular thickening diminishes the later (>24 h) proinflammatory properties of *S. pneumoniae*

Next, we tested whether exposure to conditioned medium altered the proinflammatory properties of the strains after thickening. We incubated mixed glia with bacteria with thickened capsules (preincubated for 24 hours with mixed glia) (Fig. 3A). Both bacterial strains tested (51113 (8) and 807.13 (1)) caused weaker cytokine release (TNF-α and IL-6) once their capsules thickened (Fig. 3B, C; identical CFUs). Since the proinflammatory properties of *S. pneumoniae* are enhanced by PLY(23), we also tested PLY production. The conditioned medium significantly decreased the expression of PLY, which was stronger in the capsule-thickening strain 51113 than in the nonthickening strain 50943 as a control (Fig. 3D, Supplementary Fig. S4).

**Figure 3.**
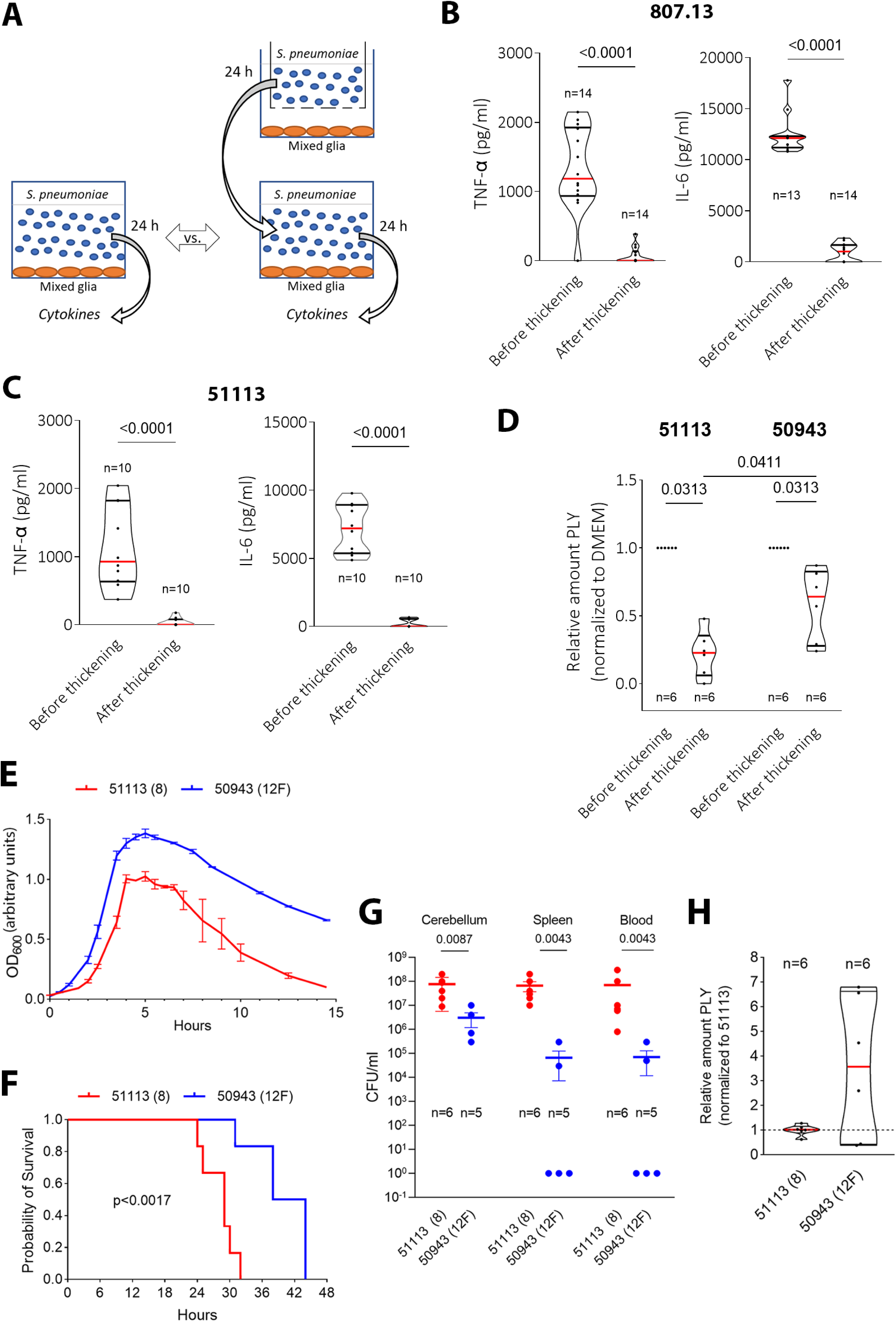
Changes in the proinflammatory, PLY production properties, and clearance of strains after capsule thickening. A. Diagram of the preconditioning of pneumococci with mixed glia and subsequent re-exposure of naïve glial cells to them for analysis of the bacterial inflammatory effect after capsular thickening. B, C. The host cell-induced increase in the thickness of the pneumococcal capsule significantly decreases the potential of serotype 1 (strain 807.13) (B) and serotype 8 (strain 51113) (C) to induce glia-dependent release of TNF-α and IL-6 (incubation with an equivalent amount of 10^7 CFU/ml pneumococci). D. Diminished amount of PLY in the supernatants of the 51113 and 50943 strains after 24 h of incubation in conditioned medium (slot blot analysis, normalized to DMEM; see Supplementary Fig. S4 as well), which is greater for the capsule-thickening strain. Each point (n) represents a biological replicate. E. Comparison of the growth curves of the 51113 and 50943 strains in BHI, demonstrating faster multiplication and higher maximum OD for strain 50943. F. Injection of 2.10^5 CFU of bacteria in the subarachnoid space produced lethal meningitis with both strains, but the progress to the endpoint is slower for the nonthickening strain (log-rank (Mantel-Cox) test). Despite the slower progression of symptoms, the 50943 strain animals also reach humane endpoint criteria. G. Number of bacteria (CFU/ml) in the cerebellum, spleen, and blood of meningitis animals at the humane endpoint experimental stop, demonstrating a continuously lower number of live pneumococci, especially in peripheral nonneural tissues of the nonthickening strain 50943. H. Comparison of the total amount of PLY (total of released and contained in bacteria) in both strains tested in DMEM (slot blot analysis) (each point (n) represents a biological replicate). For comparison of two groups, Mann-Whitney U-test was used and p-value was stated only if significant. All violin plots represent the median (red) and quartiles (black). The animal values represent the means ± SEMs of three independent experiments (E), six 51113 and five 50943 animals (F, G). In (G), no symbol for 0 CFU is shown due to the logarithmic scale.

### Capsular thickening protects bacteria against elimination and impairs the clinical course

We compared the resistance of the thickening 51113 strain and the nonthickening 50943 strain to elimination and disease progress in mice. When grown in BHI medium, 50943 bacteria multiplied faster than 51113 bacteria and reached a greater OD_600,_ indicating greater bacterial numbers (Fig. 3E). When injected in identical numbers (10^5 CFU) in the subarachnoid brain space, both strains developed lethal meningitis (the pneumococcal meningitis model in C57BL/6JRj mice is always lethal), with slower progress in strain 50943 (Fig. 3F). Compared with the bacterial growth (colony-forming units (CFUs)) of bacteria from the brain, blood and spleen, the nonthickening strain demonstrated a diminished ability for hematogenic dissemination and spleen colonization (Fig. 3G), but the number of live bacteria in the brain was lower (Fig. 3G). Initially, the expression of PLY in both strains was equivalent (Fig. 3H, Supplementary Fig. S4).

### Lack of capsular thickening in the presence of fucose and hypoxia

The nonthickening 12F strain contains a fucose residue in its capsule(30); therefore, we incubated it in conditioned medium in the presence of fucose but did not observe capsular thickening (Supplementary Fig. S5A), excluding its metabolic effect. Hypoxia stimulates capsular thickening in *S. pneumoniae*(31), and we incubated the 51113 and 50943 strains under hypoxic conditions (Supplementary Fig. S5B). 51113 demonstrated strong thickening, whereas the 12F strain remained unchanged, confirming the generally missing capsular thickness response in the latter (Supplementary Fig. S5B).

### The *cod* locus did not appear to control capsule thickening during infection

Among the South African samples, we identified a second 12F isolate (51060) that was likewise unable to thicken its capsule (Fig. 4A). In all strains studied except for 807.13 (serotype 1), the presence of both normal and inverted *cod* loci at a 1:1 ratio was observed by PCR (Fig. 4B). The lack of a *cod* locus in strain 807.13 was also confirmed by extended PCR (Supplementary Fig. S6).

**Figure 4.**
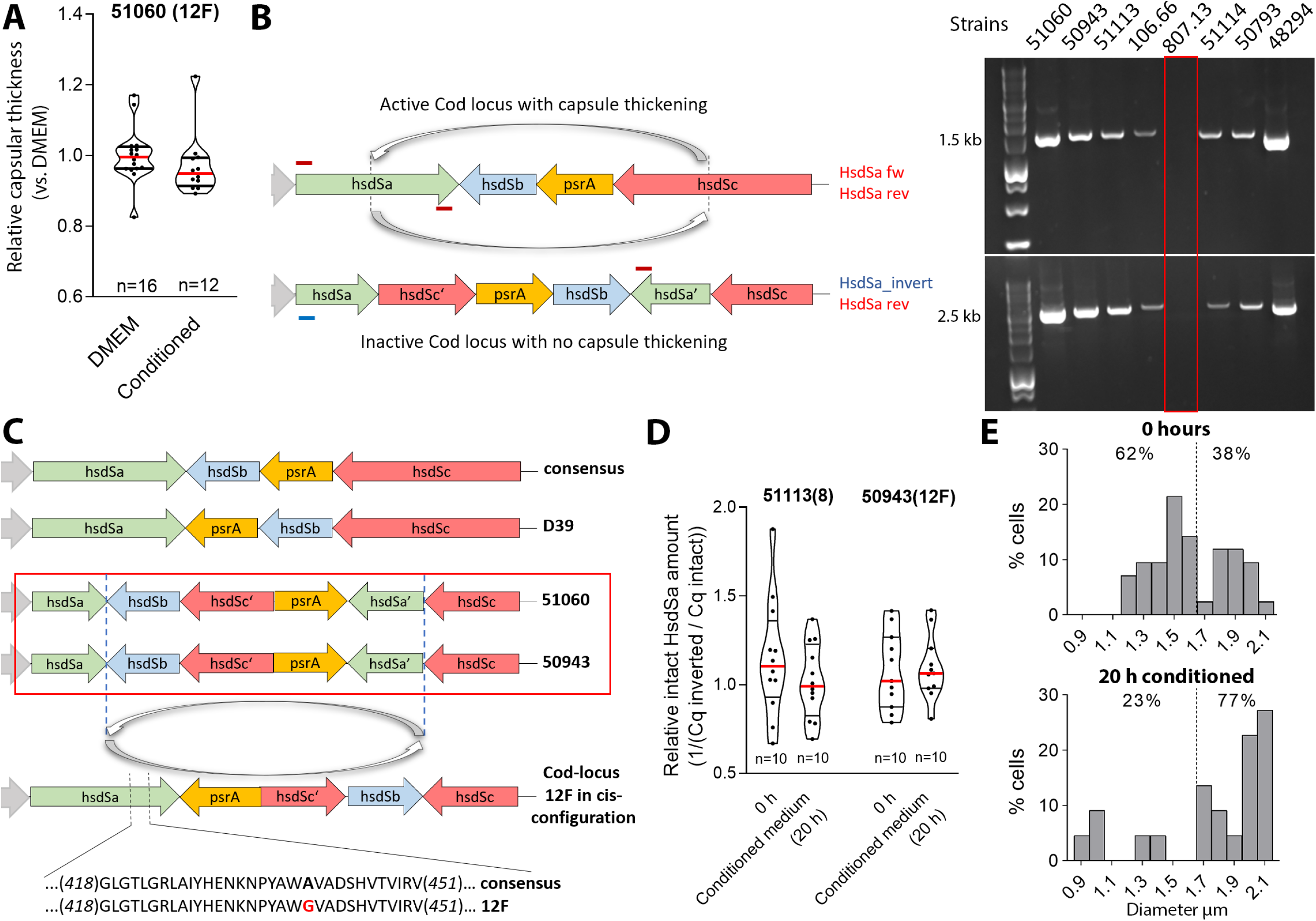
Genetic status of the *cod* locus. A. Defect in capsular thickness increase for another serotype 12F isolate from South Africa, 51060 (South Africa). Each symbol represents an image with at least four bacteria, and each experiment is performed in duplicate three times on three different days, and the data are pooled (n – number of fields). All violin plots represent the median (red) and quartiles (black). B. Diagram of the normal/inverted *cod* locus and the PCR primer position for various configuration detection methods. PCR analysis reveals the presence of both cis-and trans-cod loci in all strains except 807.13, which nevertheless demonstrates the capsule-thickening phenotype. C. Comparative diagram of the consensus *cod* locus, the D39 *cod* locus (as deposited in NCBI) and our genome sequencing data for 51060 and 50943, demonstrating an identical configuration with additional internal inversion, which does not affect the proper functioning of the sequence specificity protein hsdSa. A single point mutation at pos. 439 leads to the replacement of adenosine with guanosine, which we do not consider critical for the function of hsdSa. D. Unchanged proportions of the cis/trans *cod* loci configurations in capsule thickening 51113 and capsule nonthickening 50943 after 20 h of incubation in glia-conditioned medium (qPCR). All violin plots represent the median (red) and quartiles (black). Each point represents a biological replicate (n). E. Changes in the distribution of various capsule thickness populations in the 51113 strain after 20 h of incubation in glia-conditioned medium, which does not correspond to the unchanged 50/50 distributions of the cis- and trans-*cod* locus populations, indicating that the *cod* locus was not associated with the observed changes in capsular thickness.

Detailed genome analysis of both 12F strains revealed an intact capsular promoter (versus the laboratory strain D39; D39 also thickened its capsule in glia-conditioned medium (Supplementary Fig. S5C)). In the *cod* locus, the protein sequences of the DNA endonuclease (HsdR) and the methyltransferase (HsdM) remained unchanged (sequencing). An internal inversion of the HsdSb/HsdSc fragments, relevant to the inversion-dependent switch-off of the DNA-recognizing HsdSa, was observed, but without frameshifting HsdSa (Fig. 4C). A substitution of alanine to glycine at position 439 in HsdSa in both 12F strains was present (Fig. 4C) but unlikely to alter the methylation properties. Cis/trans *cod* locus variants remained at a similar 1:1 ratio (qPCR) before and after exposure to conditioned medium (Fig. 4D), not explaining the excessive number of bacteria with thickened capsules (>80%) in the 51113 strain at 20 h (Fig. 4E). None of the genetic configurations showed a phenotype-consistent *cod* locus presence/lack in the strains studied to explain thickening (or its lack) (Supplementary Fig. S6).

### Genetic comparison between thickening and nonthickening strains

We compared the genomes of D39 with those of the nonthickening 12F strains (50943 and 51060) (Table 1). All differing genes (>50 bp) were further compared with those of all the other thickening strains, outlining genotypes that are consistent with thickening properties (Table 1). The capsular operon promoter control sequences (preceding the core promoter), the *SPD_0642* gene (hypothetical protein, presumably a sodium-dependent transporter as determined by structural prediction), and a sequence upstream of *SPD_0994* showed consistent differences.

**Table 1.**
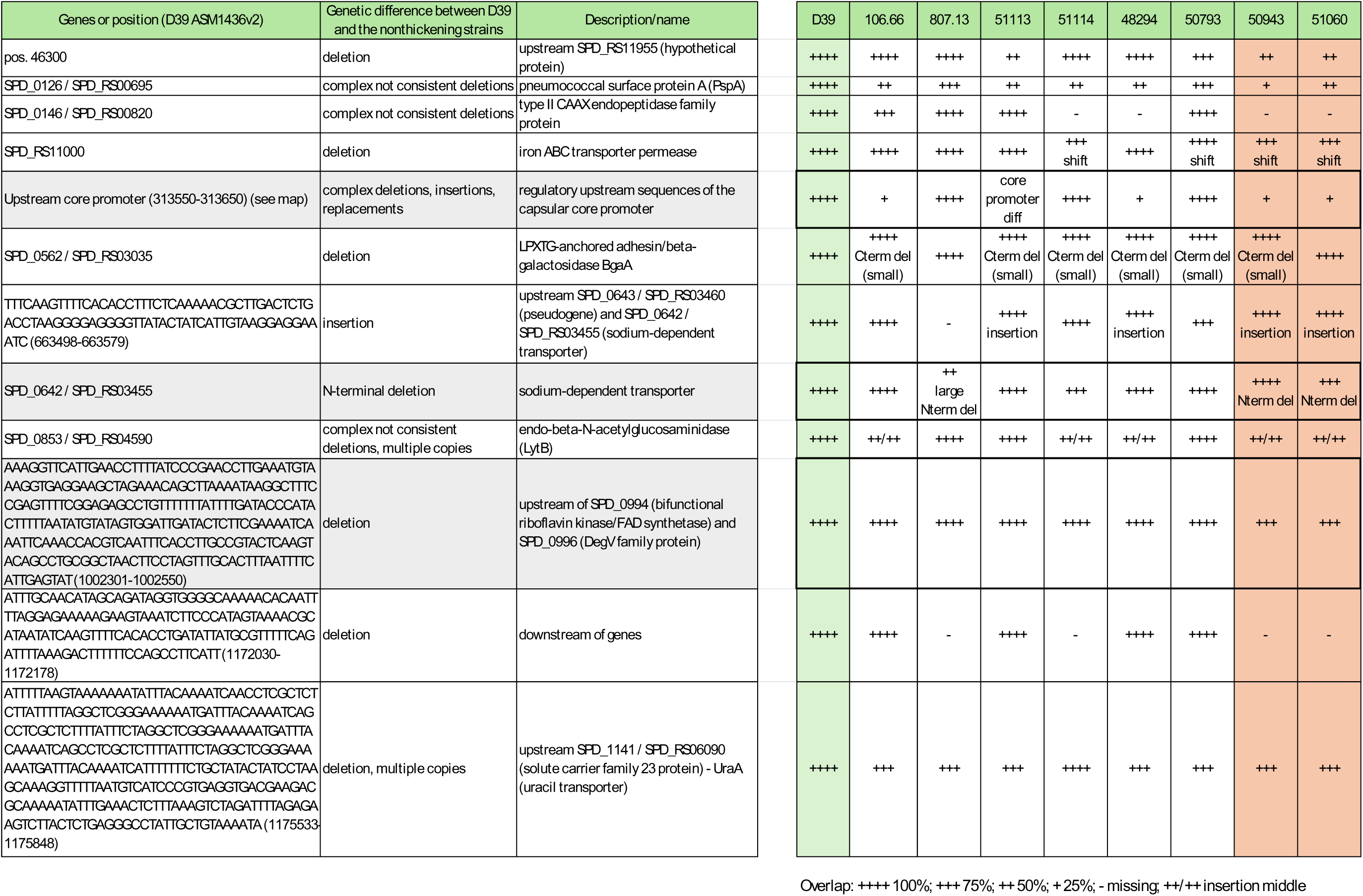
Genome comparison between thickening and nonthickening strains.

We also manually picked >500 colonies of nonthickening phenotype colonies (3 dishes) from the Mariner-transposon random genome knockout library (D39 background), pulled them together and sequenced them via Illumina next-generation sequencing (NGS) (Supplementary data D1). Here, hits in the prepromoter control sequence and in *SPD_0642* were confirmed.

### Genetic differences in capsular promoter regulatory domains

We mapped the 200 bp capsule regulatory sequence upstream of the capsular core promoter, which contains the 37-CE sequence (interacting with the capsule-altering transcription factors SpxR and CpsR) and additional sequences that interact with the regulators MalR, FabT and again CpsR (Fig. 5A)(19). No consistent differences in the capsular core promoter consistent with the thickening phenotype were observed (Fig. 5B). In the precapsular control sequence, we observed two distinct patterns. The first, observed in D39, 807.13, 50793, and 51114, and both nonthickening 12F strains, there is an existing capsular promoter regulatory domain, as shown in Fig. 5A (Fig. 5C), with consistent differences between the thickening and nonthickening strains (Fig. 5C, red outline). In three other strains, 106.66, 48294 and 51113, no prepromoter sequence consistent with Fig. 5A was detected (Fig. 5D). To verify the functional importance of these differences, we constructed mutants of D39 with a completely missing prepromoter control sequence (Fig. 5E) or replaced with the 50943 (nonthickening) strain (Fig. 5E). Challenge of these mutants with glia-conditioned medium revealed that bacteria whose 200 bp regulatory sequence was completely missing (Δ37-CEreg) still had thickened capsules, albeit slightly less (Fig. 5F). Replacing the regulatory sequence in D39 with the 12F-specific prepromoter sequence (37-CEreg 12F) resulted in weaker thickening than D39 (Fig. 5F). Thus, when missing, the prepromoter control sequence had little effect on thickening, but when present, the 12F-specific sequence diminished capsular thickening.

**Figure 5.**
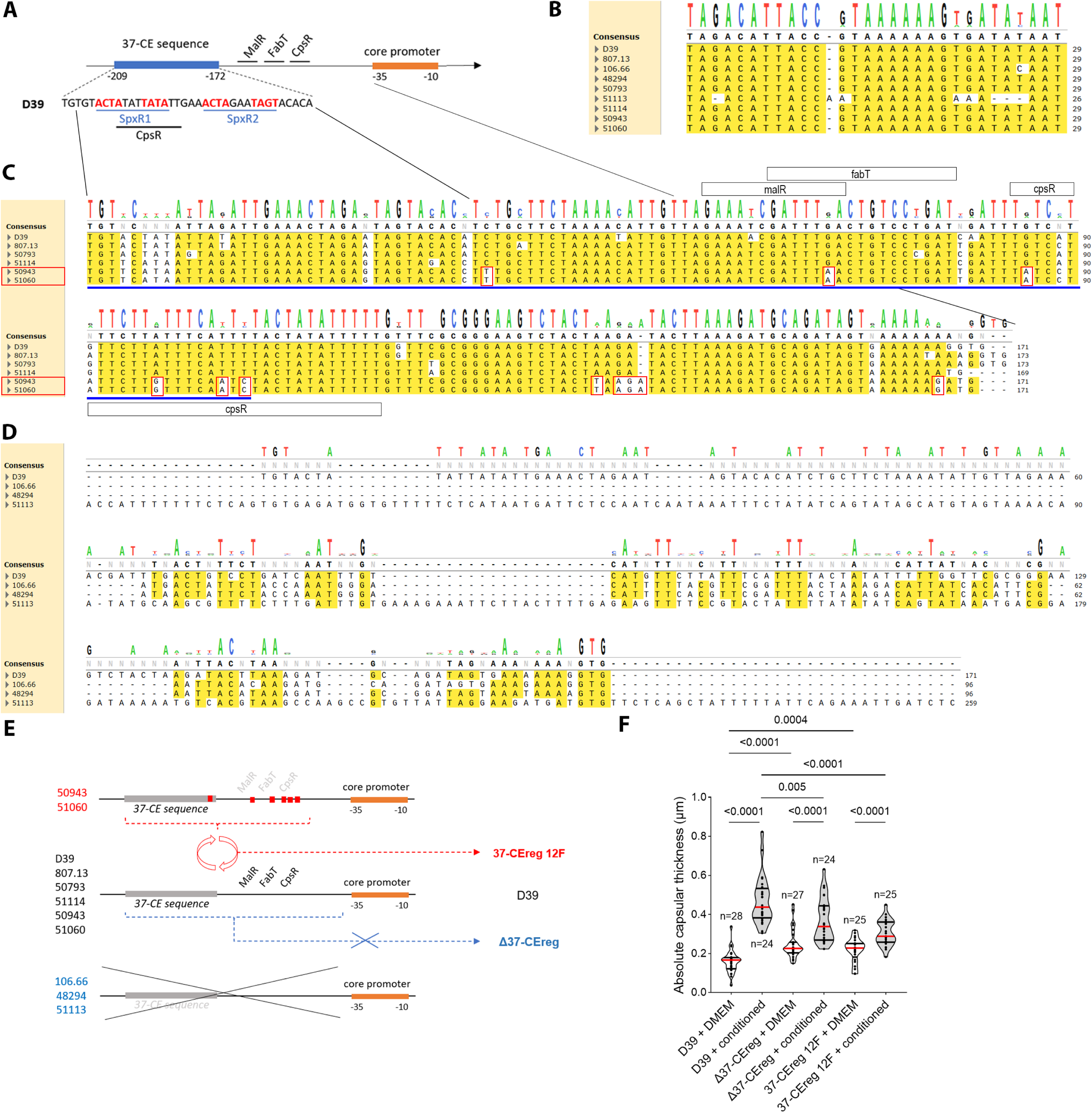
Comparison and thickening role of capsular promoter regulatory domains. A. Schematic diagram of the regulatory domain positioned approximately 200 bp upstream of the capsular core promoter. The 37-CE sequence is also shown in full length with corresponding transcription factor docking domains. B. Comparison of the exact sequences of core promoters in all tested strains with only minimal differences, which is not consistent with the thickening phenotype. C. Alignment (Clustal Omega) of the sequences of the regulatory domains of strains D39, 807.13, 50793, 51114, 50943 and 51060, demonstrating high homology with point mutations in the docking sequences for the transcriptional regulators malR, fabT and cpsR in the 12F strains (outlined in red) versus all other strains. D. Lack of homology in the regulatory domains of strains 106.66, 48294 and 51113 versus D39. E. Schematic representation of the generated mutants in the regulatory domain of the capsular promoter: Δ37-CEreg – completely removed regulatory domains upstream of the core promoter; 37-CEreg 12F – replacement of 115 bp of D39 with the corresponding sequence from the nonthickening 12F strains. The regions of discrepancy between thickening and nonthickening strains are schematically presented in red. F. When exposed to glia-conditioned medium, both Δ37-CEreg and 37-CEreg 12F mutants still thickened their capsules, but less than native D39. Each symbol represents one image containing at least four bacteria. Each experiment is performed in duplicate three times on three different days, and the data are pooled. Violin plots represent the median (red line) and quartiles (black line). All strains are compared via Kruskal–Wallis with Dunn’s multiple comparisons, and the p value is presented if significant.

### *SPD_0642* is important for capsular thickening

Blast-based sequence prediction and AlphaFold 3.0 analysis suggested sodium-dependent transporter properties for *SPD_0642*. The 12F strains presented a truncated N-terminus (47 amino acids due to a frame shift with a point mutation) (Fig. 6A, Supplementary Fig. S7). One of the thickening strains, 807.13, had greater truncation of the N-terminus with preserved thickening, and one strain, 51114, had a small deletion in the middle. To study gene function, we generated a truncated N-terminus (ΔNterm SPD_0642) mutant in the D39 background (Fig. 6A). It could not increase its thickness upon exposure to glia-conditioned medium (Fig. 6B). AlphaFold 3.0 3D reconstruction revealed a transmembrane protein configuration with an N-terminus extending outside the protein barrel (Fig. 6C). C57BL/6JRj mice infected subarachnoidally with identical numbers of wild-type D39 (10^5 CFU), ΔNterm SPD_0642 and 37-CEreg 12F bacteria demonstrated slightly improved survival following infection with N-truncated *SPD_0642* (Fig. 6D). Behavioral scoring, however, showed significantly diminished symptom score in ΔNterm SPD_0642 (Fig. 6E). The bacterial loads in the brain, blood and spleen remained identical among groups (Fig. 6F). None of the mutants demonstrated impaired multiplication (Supplementary Fig. S8) (32, 33). Homology analysis via OrthoDB among various bacterial classes revealed homologs of *SPD_0642* in multiple Gram-positive and Gram-negative pathogens, with variable levels of homology (Fig. 6G). The regulatory sequence upstream of *SPD_0994* did not affect capsular thickening (Fig. 6H).

**Figure 6.**
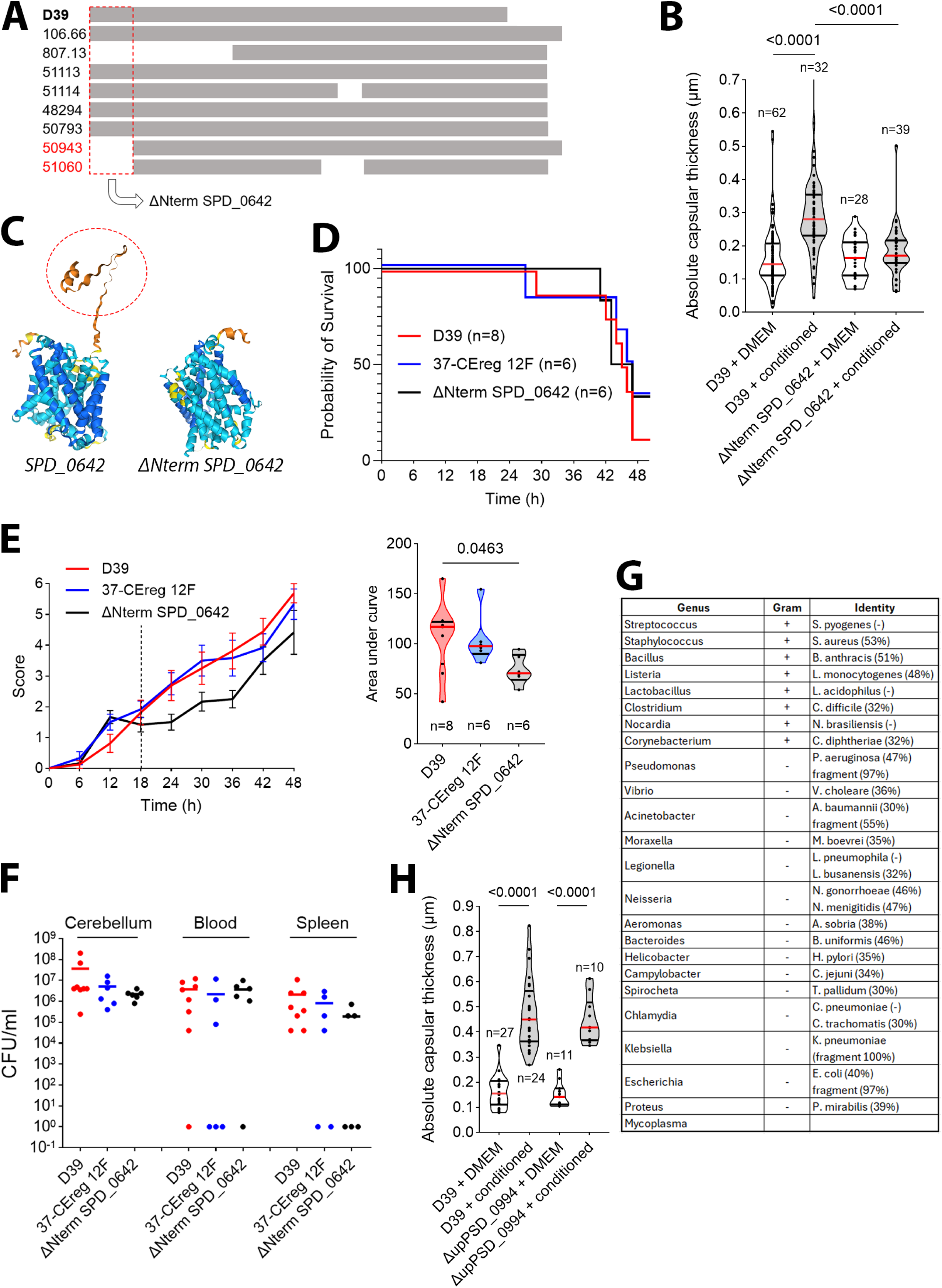
Structure and effects of *SPD_0642*. A. Schematic diagram of the *SPD_0642* gene and the deletion of the N-terminal domain in the 12F strains (outlined in red). Only 807.13 of all the thickening strains contain a larger N-terminal trimmed fragment than 12F. B. The N-terminal truncated mutant (ΔNterm SPD_0642) still forms a capsule but lacks capsular thickening in glia-conditioned medium. C. AlphaFold 3.0 3D modeling shows a well-packed transmembrane configuration with the N-terminus outside its core and high motility. The N-terminal free chain is fully absent in the 12F mutants. D. Survival curves of C57BL/6JRj mice injected in the subarachnoid brain space with 10^5 CFU of each presented bacterium. Curves were analysed with log-rank (Mantel-Cox) test. E. Behavioral scoring of all mice injected with D39 and mutant strains, showing significantly reduced clinical scores in ΔNterm SPD_0642 (area under curve, right graph). On the left graph, all values indicate means ± SEMs. F. CFU/ml in the brain (cerebellum), blood and spleen of meningitis animals at the time of sacrifice, demonstrating similar bacterial numbers. G. Levels of identity of *SPD_0642* in clinically relevant representatives. H. Preserved capsular thickening by glia-conditioned medium in D39 pneumococci with removed noncoding sequence upstream *SPD_0994*. In the violin plots, each symbol represents one image containing at least four bacteria. Each experiment is performed in duplicate three times on three different days, and the data are pooled (n - number of counted fields). Violin plots represent the median (red line) and quartiles (black line). All strains are compared via Kruskal–Wallis with Dunn’s multiple comparisons, and the p value is presented if significant.

## Discussion

We show that *S. pneumoniae* dynamically changes its capsular thickness in response to host tissue (noninflammatory and nonnutritional effects) and identify *SPD_0642* (a predicted sodium-dependent transporter) as a previously unrecognized capsule modulator gene. Capsular thickening was tissue-specific - most pronounced upon exposure to brain tissue, moderate in serum, and absent on epithelial surfaces. Once established, this thickening correlated with reduced late neuroinflammatory capacity, consistent with lower PLY expression. Nonthickening serotypes exhibited reduced dissemination beyond the brain and slower/milder disease progression yet remained lethal when directly introduced in the subarachnoid brain space. Genetic analysis excluded a dependence on *cod* locus configuration but indicated a partial contribution of the upstream regulatory sequence of the capsular operon and the involvement of *SPD_0642*.

Bacterial capsules protect pathogens from phagocytosis, complement attack, and immune clearance but also impede antibiotic penetration, fostering tolerance and potential resistance (34). Elimination of capsular genes often improves spontaneous bacterial clearance by the host (35). The epidemiological persistence of nonencapsulated *S. pneumoniae* lineages underscores some niche context-dependent advantages (36) (37). The impact of capsular thickness, however, remains underexplored. *Acinetobacter baumannii* (38, 39) and *S. pneumoniae* (40, 41) benefit from thicker capsules to reduce mucus binding and improve cell adhesion. Previously, we reported that *S. pneumoniae* with a thicker capsule causes more severe meningitis(12). Only moderate capsular thickness improves upper respiratory tract colonization by pneumococcus(42). On the epithelium, the pneumococcal capsule becomes thinner(43). Thicker capsules correlate with greater virulence in *Klebsiella* (44), whereas thinner capsule beta-hemolytic *Streptococci* demonstrate greater virulence and stronger blood‒brain barrier penetration(45). Capsular thickening is observed in *Streptococcus suis* when incubated in a metabolic chamber in the peritoneum of rodents, with unclear disease importance(46). Capsular thickness correlated with neutrophil extracellular traps in lung infections by *S. pneumoniae*(47). The thickening of the capsule of *Ruminococcus gnavus* in the intestine leads to milder inflammation and better symbiotic coexistence between bacteria and the host(48). In *N. meningitidis*, another neurotropic bacterium, capsular thickening is enhanced by fever during infection(49). While capsules improve resistance against the immune system, they can also impair barrier penetration and are energetically demanding(50) (43).

Phase variation via *cod* locus inversion generates capsule-rich and capsule-poor subpopulations(51). Capsular thickness changes due to nutritional differences, such as source of carbon, oxygen level, and sugar type(52). Our findings show that capsular thickening can occur independently of the *cod* locus inversion, implicating additional regulation. Nutritional parameters remained constant across experiments, ruling out metabolic triggers. Instead, the inducing factor appears soluble, signaling in nature, and noninflammatory, as it was secreted even by resting glial cells.

StkP (with its cognate phosphatase PhpP) is a central serine/threonine signaling module in pneumococcus that links envelope/cell-division physiology to transcriptional and metabolic programs; it can influence capsule operon expression/capsule output(53). Phosphorylation of the critical capsular gene CpsD diminishes capsular expression, further suggesting a possible kinase/phosphatase-based regulatory switch(54).

In the opportunistic *Cryptococcus neoformans,* various factors (low CO_2_, low glucose and nitrogen, stress and low iron (all environmental or nutritional stress conditions)) lead to capsular thickness changes, altering pathogenicity (55).

Capsular thickening is described in Gram-negative bacteria (*Klebsiella pneumoniae*) by the factors RmpA2 and RcsB(56). The polypeptide antibiotic polymyxin B and the antibacterial lactoferrin alone can enhance the protective capsule of *K. pneumoniae*(*57*). Similarly, sublytic antibiotic treatment in *Acinetobacter baumannii* causes capsular thickening and increases bacterial antibiotic resistance(58). Like *Klebsiella*, a two-component system (BfmRS) is also involved(59).

In *S. pneumoniae*, *SPD_1495* acts as a negative regulator of capsular synthesis, interacting with ComE(60, 61). Thus, molecularly, the system of capsular thickening regulation appears to often involve two-component systems – a transcriptional regulator and a sensor – suggesting the ability of environmental control. Under the control of a quorum-sensing (bacterial pheromone-dependent) system, *S. pneumoniae* also thickens its capsule, suggesting environmental control(62).

Pneumococci can sense and respond to peptides secreted from other bacteria, resulting in interspecies communication(63–65). Comparable sensing of host-derived signals is plausible. The factor responsible for thickening was active in conditioned medium but not in brain and heart homogenate medium. Possible explanations include degradation during preparation or uneven secretion and dilution. Fetal calf serum, also heat-inactivated, induced only mild thickening, indicating that heat inactivation alone does not abolish activity. Incubation with DMEM, like the effect of epithelial cells, may also lead to thinning of the capsules and thus mask brain-heart-infusion thickening.

Bacterial capsules control both the inflammatory response and phagocytosis resistance. During early infection, thickening strains triggered stronger cytokine responses (IL-6, TNF-α, CXCL2), but by 24 h, once they thickened, they became less proinflammatory. This inverse association could reflect late PLY downregulation, consistent with the known inflammatory activity of PLY(23). Disease progression in nonthickening strains was slower but ultimately fatal, confirming that thickening shapes the kinetics, not the outcome, in our infection model. We have not studied, however, functional deficits in combination with antibiotics, which can also reveal other functional effects of capsular thickening, as well as other invasion ways.

The upstream regulatory region of the capsular operon contributed to, but was not essential for, thickening. Its known control elements, such as the 37-CE sequence, bind spxR and cpsR to repress transcription(19). The remaining consensus sequence (D39) contains transcription-sensitive binding sites for spxR(19), cpsR(66), malR(19), and fabT(67), which are all expressed in all our strains. Strains (106.66, 51114 and 48294) lacking this upstream regulatory region still thickened, implying compensatory regulatory motifs. Mutations in the malR, fabT, and cpsR binding sites distinguished nonthickening 12F strains and likely disrupted cooperative regulation. Thus, capsular control appears multifactorial, with overlapping transcriptional checkpoints.

The identification of *SPD_0642* as a capsule thickening modulator was unexpected. Bioinformatic analysis predicts it as a membrane-associated transporter related to the SLC5/6 family, typically involved in amino acid or neurotransmitter transport(68) (preserved both in prokaryotes and in higher organisms). Further interactome and experimental analysis is required to clarify its exact role for the capsules(69). Truncation of its N-terminus removed a separate, relatively free moving tail, not included in the stably packed beta-sheet core of the protein (AlphaFold). Whether this impaired membrane localization of the protein is unknown, bacteria still possessed capsules, indicating partially preserved function. The 807.13 strain lacked a larger fragment of the gene and still exhibited thickening, suggesting some redundancy. In both 12F strains, the combination of both elements — a truncated *SPD_0642*, and a modified sequence upstream the capsular promoter – may alter thickening more potently than each alone. For mechanistic reasons, we tested them separately. *SPD_0642* is widespread (with variations) among many Gram-positive and Gram-negative bacterial strains. Whether its effect on capsules is similar/identical to that of *S. pneumoniae* requires further studies.

### Limitations

i. We tested a gene variation in *SPD_0642* and in the capsular prepromoter sequence on thickening separately, and we cannot exclude more complex interactions between them.
ii. The analyzed strain panel was heterogeneous, but cross-validation across multiple isolates minimized strain-specific bias.
iii. Additional modulatory genes with partial effects may exist, potentially amplifying or dampening the dominant thickening pathways.

## Conclusion

Capsules in *S. pneumoniae* are dynamically regulated defense structures, not static traits. Their plasticity modulates immune interaction, dissemination, and disease kinetics. The discovery of *SPD_0642* as an extrinsic regulator of capsule thickening uncovers a serotype-independent vulnerability that could eventually be exploited for targeted antimicrobial intervention.

## Materials and Methods

### Bacterial strains and culture

The strains used were recovered from the cerebrospinal fluid of meningitis patients in 2017 in South Africa and from nasopharyngeal samples of the Swiss Nationwide Surveillance Program (70). The South African strains were collected within the South African GERMS national laboratory-based surveillance program and kindly provided by the National Institute for Communicable Diseases (NICD) in Johannesburg. Sample collection was performed with approval from the Cantonal Ethics Commission (Bern, Switzerland) and the Human Research Ethics Committee (Medical) of the University of Witwatersrand (Johannesburg, South Africa). The relevant university and provincial ethics committees approved the GERMS-SA surveillance study (clearance nos. M140159, M081117, M021042, and M180101). The serotypes of all the strains were confirmed via the Quellung reaction. We also used the capsule deletion mutant of the D39 strain (a kind gift from Jeremy Brown). All strains with serotype information are listed in Supplementary table T1.

For long-term conservation, the strains were stored at -80°C in Protect bacterial preservers (Technical Service Consultants, Heywood, U.K.) or in 10% glycerol in brain-heart infusion broth (BHIB, Becton Dickinson and Company, le Pont-de-Claix, France). Bacterial suspensions were prepared by streaking bacteria on Columbia sheep blood agar (CSBA) plates and subsequently incubating them at 37°C and 5%CO_2_ overnight. The experimental samples were prepared from stocks as close as possible to the original, avoiding multiple passages. Multiple colonies were picked, inoculated in BHIB and grown at 37°C to OD =0.5–0.7 (mid‒to-late log phase). After three washes in 1x PBS, serial dilutions were plated on CSBA to determine the CFU/mL. For all experiments, bacteria were diluted to obtain a final concentration of 10^7 CFU/mL in Dulbecco’s modified Eagle medium (DMEM with high glutamate; Gibco, Thermo Fisher Scientific AG, Basel, Switzerland).

Hypoxia was initiated in an anaerobic jar via anaerobic culture fast packs (Sigma‒Aldrich Chemie GmbH, Taufkirchen, Germany) and verified via Anaerotest strips (Sigma), which contained <0.1% oxygen.

For the defensin experiments, human β-defensin 3 (hBD3, Sigma) was used.

### Genetic manipulation of D39

Bacteria were mutated as described previously(71). In brief, primers (Tm=60°C; primers in the table) corresponding to the flanking regions of the gene of interest (approximately 1,000 bp upstream (primers F1 and R1) and downstream (primers F2 and R2) of the region of interest) were amplified. The primers directly neighboring the region of interest contained spectinomycin overhang sequences. PCR amplification was performed by high-fidelity Q5 polymerase (New England Biolabs) for 30 cycles (98°C for 10 s, 60°C for 30 s, and 72°C for 60 s), including amplification of the spectinomycin resistance gene. Following gel extraction from low-melting agarose (stained with SybrGreen (Thermo Fisher) under blue light) and gel extraction (MinElute gel extraction kit (Qiagen GmbH, Hilden, Germany), the three fragments (the amplified spectinomycin gene and both flanking regions, 50 ng each) were annealed without primers in the presence of Q5 polymerase for 25 cycles (98°C for 30 s, 52°C for 30 s, and 72°C for 90 s). Specifically, 2 µl of the 50 µl reaction mixture was transferred to a new PCR amplification mixture containing both end primers of the annealed 3-fragment sequence (3 kb product expected) and run for 35 cycles. Following gel extraction of the amplified fragment, freshly grown D39 (OD=0.1, either in brain heart infusion broth (BHIB) or Todd-Hewitt broth (THY) with oxyrase (1:200) (all from Sigma)) was incubated with 0.02 volumes (20 µl) of 1 N NaOH, 25 µl of 8% BSA, 1 µl of 1 M CaCl_2_ (all from Sigma), 1.5 µl of 350 ng/µl CSP-1 (competence-stimulating peptide-1 (AnaSpec Inc., Fremont, CA, USA), and 350 µg/ml) for exactly 15 min at 37°C. Half of the gel-extracted DNA was incubated with the bacteria for 45 min and subsequently seeded on spectinomycin (Sigma)- containing (200 µg/ml) blood agar plates. The resulting colonies were further grown in spectinomycin-containing broth, washed and frozen in 12.5% glycerol (Sigma)-containing broth or sent for genome sequencing (Microsynth AG, Balgach, Germany) and verification of the replacement of the gene with the spectinomycin gene.

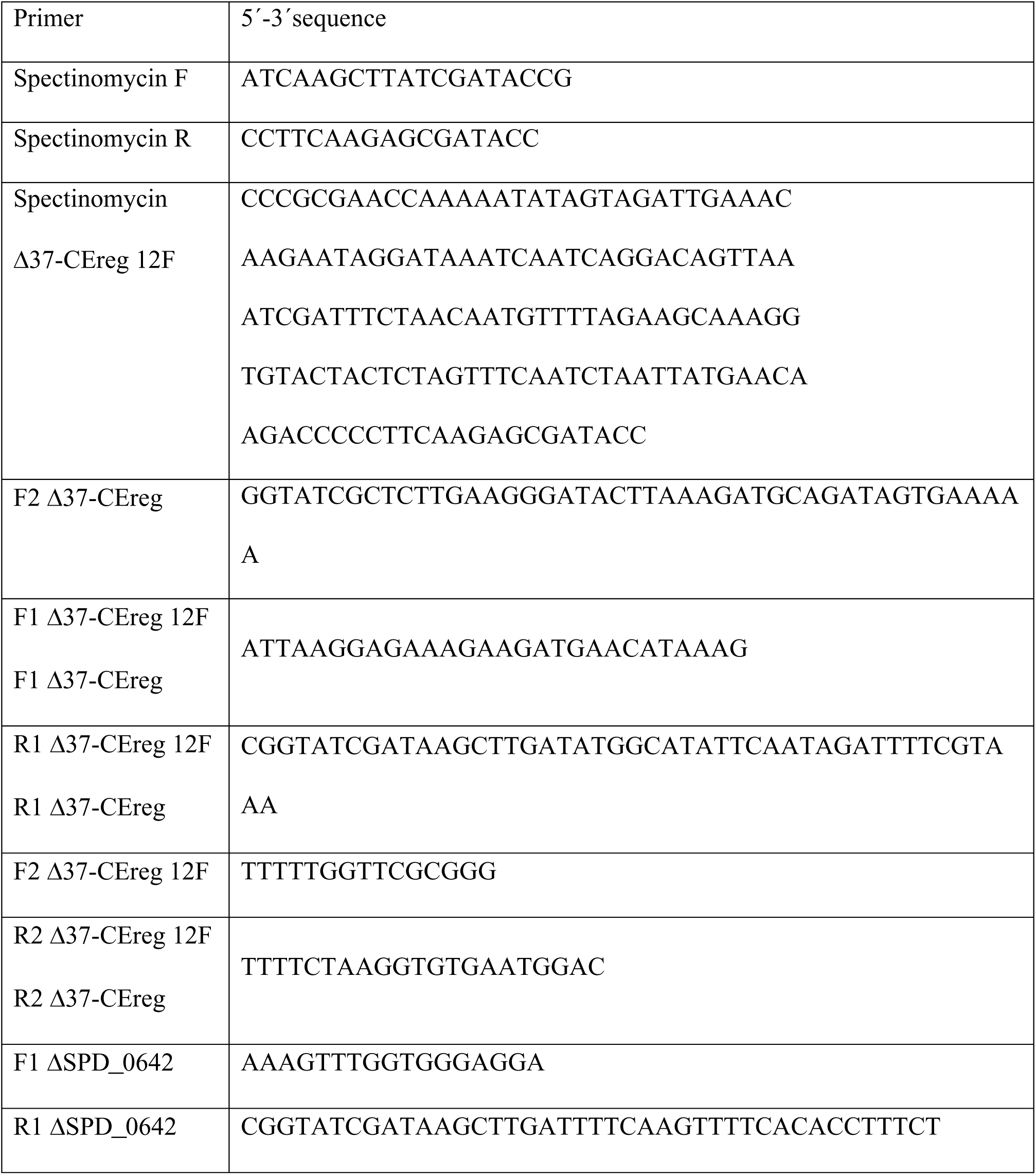
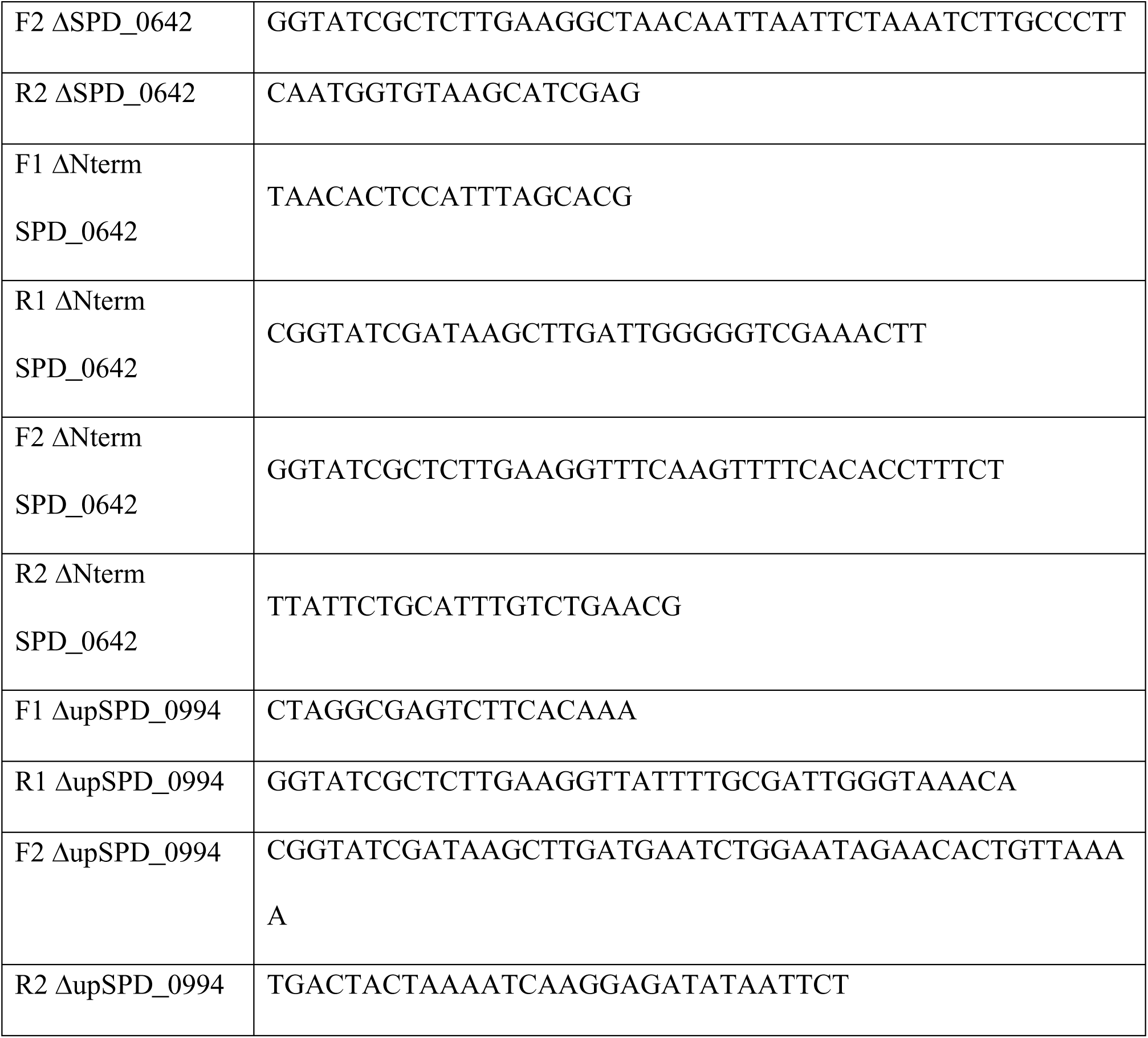

### Cell cultures and glia-conditioned medium preparation

Primary mouse astrocytes and microglia were prepared from the cortices of newborn C57BL/6JRj mice (postnatal day (PD) 3-4) as mixed cultures. Briefly, the cortices of newborn mice were dissociated into cell suspensions and plated in cell culture flasks (75 cm^2^) (Sarstedt AG, Nuembrecht, Germany) coated with poly-L-ornithine (PLO, Sigma). The growth medium, DMEM, was supplemented with 10% inactivated FCS (PAN Biotech, Aidenbach, Germany) and 1% penicillin/streptomycin (Gibco). Eleven to fourteen days after preparation, the cells were ready for use in the experimental procedures.

Glia-conditioned medium was prepared from mixed glial cultures after day in vitro 14 as follows: adherent cells were washed 4 times extensively with PBS to remove all antibiotic and serum rests and incubated for 24 h in antibiotic- and serum-free DMEM. It was changed again with antibiotic- and serum-free DMEM for 24-36 h and harvested. Following centrifugation for 7 min at 3,000 g for removal of all debris, glia-conditioned medium was used either immediately or stored frozen at -20°C.

### Media fractionation and treatments

Amicon® Ultra Centrifuge filters (Millipore, Sigma) with cutoff molecular weights of 10, 5 and 3 kDa were used to exclude all higher-molecular-weight components. The columns were extensively washed with 70% ethanol, aspirated, centrifuged (not touching the membrane) and washed 3 times with distilled water. Finally, the samples were centrifuged (20 min) to remove the excess water before adding medium.

### Cytokine measurements

Murine mixed glia were plated on PLO-coated 24-well plates (200,000 cells/well) and tested after 7 days. Before treatment, the cells were washed once with serum-free medium. The levels of TNF-α, IL-6 and CXCL2 (MIP-2) were determined as early proinflammatory responses of the mixed glia to bacterial infection. Cytokine release was measured after 24 h of incubation with live bacteria at 37°C and 5%CO_2_ in DMEM without serum. To determine the role of increased capsule size in the proinflammatory response, we cocultured bacteria in cell culture inserts (Sarstedt, 0.4 µm pore size) together with glial cultures for 24 hours. No bacteria were found outside the insert. 10^7 CFU/ml were used. Bacterial counts were determined every time prior to infection by plating serial dilutions on CSBA. Cytokines were measured with conventional sandwich ELISA against mouse TNF-α, IL-6 (BioLegend ELISA MAX, San Diego, CA, USA) and CXCL2/MIP-2 (R&D Systems, Minneapolis, MN, USA) following the manufacturer’s instructions. The absorbance was measured at 450 nm with an EL800 microplate reader (BioTek Instruments, Winooski, VT, USA).

### Capsule thickness measurement

We measured pneumococcal capsule size via the FITC-dextran exclusion assay as described previously (72). In the initial experiments, we measured total bacterial diameters, which correlated with capsular thickening on a widefield microscope, as data were presented in pixels. In the next experiments, we switched to the more precise direct measurement on a confocal microscope, subtracting negative contrast from transmission image (Supplementary Fig. S1A, B). In these later experiments, we presented the results as absolute (µm) or sometimes as relative thicknesses (versus the mock-control) depending on the aim and the clarity of presentation. Normally, thickness was measured before and after 9 and 24 hours of incubating the strains (10^7 CFU/mL) with primary mixed glia directly or with conditioned medium. In the setup without contact, the bacterial suspension was pipetted into a tissue culture insert (0.4 µm pore size, Sarstedt), which served as a barrier between bacteria and cells but enabled the exchange of soluble factors. A total of 10 µl of bacterial culture was mixed with 2 µl of FITC-dextran (2,000 kDa, Sigma; 10 mg/ml in water) and then pipetted onto a microscope slide. A coverslip was used to trap the liquid. The slides were viewed on a Zeiss Axio Imager M1 (Carl Zeiss GmbH, Jena, Germany) fluorescence microscope with a 100X objective and a FITC filter or on a Zeiss LSM880 with a 63x objective, 2x zoom and 0.05–0.2% power with a 488 nm argon laser. Because of system differences, in some experiments, data were presented as average pixel area/bacterium (wide field), normalized (relative) area (confocal) or direct thickness measurements (confocal). Each experiment was performed in duplicate and repeated three times on 3 different days, adding up to 30 images/strain per time point (9 and 24 hours).

### Quantitative PCR

PCR genome mapping of the cod locus was performed via 40 cycles of PCR master mix (MedChemExpress LLC, NJ, USA) (20 s at 96°C, 30 s at 56°C, and 90 s at 72°C). Quantitative PCR was performed with identical cycle parameters using SybrGreen master mix with high ROX (MedChemExpress LLC) on a QuantStudio 6 Pro (Thermo Fisher). Curves were analyzed by Design & Analysis Software (ver. 2.6.0, Thermo Fisher). The following primers were used:

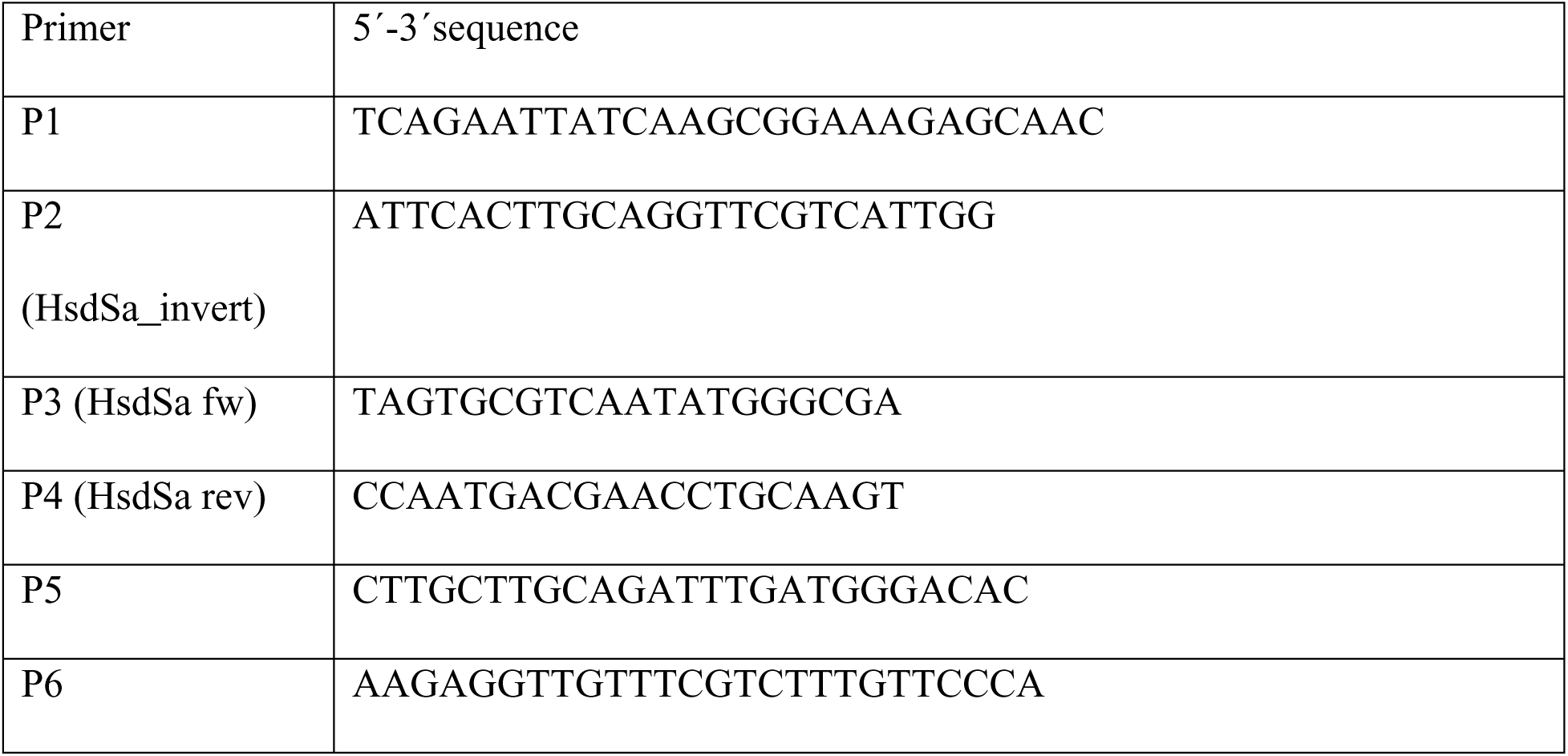

### Protein biochemistry

Bacterial samples were collected in RIPA buffer, heated to 95°C with protease inhibitor mixture (Roche Diagnostics GmbH, Mannheim, Germany), and further analyzed by slot blotting. The protein amount was initially roughly assessed via a BCA protein assay kit (Boster Biological Technology, Pleasanton, CA, USA) and was precisely adjusted thereafter via Coomassie staining of the PVDF membranes (Amersham plc, Amersham, UK). For the slot blots, samples were aspirated through the membrane and washed with PBS-T (0.05% Tween-20 (Sigma)). Thereafter, the membranes were blocked with 3% skim milk (Sigma) in PBS-T and incubated with primary antibodies against PLY (1:200, mouse monoclonal (PLY-4), ab71810, Abcam plc, Cambridge, UK) and HRP-tagged secondary goat anti-mouse antibodies (1:2,000, Sigma). Finally, the membranes were visualized with an enhanced chemiluminescence (ECL) substrate (Immobilon Western, Merck Millipore, Burlington, MA, USA).

### Animal experimentation and tissue preparation

#### Induction of meningitis

All animal experiments were approved by the Cantonal Animal Protection Commission in Bern (BE103/2020 and BE91/2025). We used 8-week-old C57BL/6JRj (male and female) mice (DBMR, Department for Biomedical Research, University of Bern). Meningitis was induced by injecting 2.10^5 CFU of strains 51113 and 50943 in 10 µl in the subarachnoid space at the following coordinates: 2 mm anterior from the bregma, midline, and 4 mm deep(73). In the experiments with thickening-deficient strain mutants, 10^5 CFU was used to achieve a milder disease course and better discrimination of disease progress. Additionally, compared with the initial 51113 and 50943 experiments, we applied more aggressive rehydration and glucose application strategy in accordance with the requirements of our ethical commission. The animals were anesthetized with 3% isoflurane and maintained with 1.5% isoflurane (Baxter AG, Opfikon, Switzerland). Before the incision, the skin was infiltrated with a mixture of lidocaine/bupivacaine. All animals were kept on a heating pad during surgery and housed in a 37°C incubator immediately after surgery. Food and water with easy access (DietGel Criticare, ClearH2O, Westbrook, ME, USA) were provided. The animals were monitored following a scoresheet every 3 h with parameters such as activity, coat, status of the orbita (“orbital tightening”) and neurologic status.

#### Bacterial isolation

Twenty-four hours after induction, the mice were sacrificed by decapitation under anesthesia. The brains were harvested and washed in 1 ml of sterile PBS. The samples were centrifuged at 700 rcf for 5 min to remove all eukaryotic cells. The remaining samples were used for bacterial count and capsular thickness assessment (when performed). For bacterial CFU assessment, half of the cerebellum, a spleen and 100 µl of blood were homogenized/diluted serially and seeded on blood agar.

#### Intranasal colonization

Ten microlitres of bacteria, which were grown to OD=0.6, washed 3 times with PBS, and dissolved in 10 times lower volumes of PBS were instillated into each nostril by passive inhalation in isoflurane-anesthetized animals. The animals were allowed to recover and did not show any signs of altered health. Twenty-four hours later, the animals were sacrificed. A 1 ml syringe containing 100 µl of sterile PBS with a sterile peripheral venous catheter of the smallest size was used to wash the nasal cavity. The catheter was inserted through the trachea until it reached the nasal cavity, which was verified by the appearance of water through the nostrils when the syringe was pressed. PBS was collected and centrifuged at 500 × g to remove red blood cells or epithelial cells, and the bacterial capsular thickness of the pneumococci was evaluated. Pneumococci were identified morphologically; their number and carriage were determined via CSBA.

#### Scoring

Animal scoring following the experiments was performed according to the following semiquantitative scale: activity: normal to mildly diminished, reactive to stimulus – 0; mildly diminished – 1; moderately diminished, no active exploration, hiding – 2; strongly diminished, coma or not reactive to stimuli – 3. Coat: normal – 0; general lack of grooming – 1; piloerection – 2; posture and face expression: normal, straight back, eyes open, ears in forward-facing position – 0; mildly hunched back, orbital tightening – 2; strongly hunched back, strong orbital tightening, ears in backward-facing position – 3. Neurologic status: normal – 0; ataxia – 3. Any individual score of 3 was considered an interruption criterion (humane endpoint).

#### Activity measurement

Animals were placed in a standard activity box (50 × 50 cm), and automated camera-based tracking (Stoelting Co., Wood Dale, IL, USA) was performed for a period of 5 min. Several parameters, such as total travel distance, mean and peak velocity, and duration of activity, were automatically generated.

### DNA extraction, whole-genome sequencing, Mariner transposon screening and bioinformatics

Bacteria were streaked on CSBA from protected bacterial preserves and incubated overnight at 37°C and 5% CO_2_. Genomic DNA was extracted from a full CSBA plate of bacteria via the QIAamp DNA Mini Kit (Qiagen) according to the manufacturer’s protocol. Next-generation sequencing (NGS) was performed via the Illumina Nextera DNA Flex assay (V2, 2 × 150 bp, 300 cycles) on an Illumina MiSeq benchtop sequencer (Illumina) according to the manufacturer’s protocols. The sequencing data were processed and assembled via the Shovill pipeline (https://github.com/tseemann/shovill) with the –trim option and default parameters (v0.9.0) (Microsynth). Compiled and annotated sequences (partially available online already annotated, partially performed by tools at www.kbase.us) of all the strains were compared via Mauve (multiple genome alignment) software (the Darling laboratory, University of Technology, Sydney; https://darlinglab.org/mauve/mauve.html). For the experiments with the Mariner transposon library, we followed a previously described protocol(71). In brief, a library pool of Mariner transposon random gene knockout pneumococci was incubated with glia-conditioned medium for 8 h, followed by seeding on THY broth (Sigma) without blood supplementation for observation of transparent/opaque colonies(74). After at least 500 colonies were picked from three independent experiments, the colonies were pooled and allowed to grow until the OD reached 0.3. Following lysis and DNA extraction (Qiagen), the genomic DNA was digested with Mme I (NEB) to release the genomic fragments next to the transposon. Following CIP (NEB) treatment, the barcode adapters were annealed, and the fragments were amplified via 22 cycles of Q5 high-fidelity polymerase. Finally, the fragments were separated and extracted on a SybrGreen-stained gel. After gel extraction (Qiagen), the sequences were pooled together with other experimental samples and sequenced through an NGS pipeline (Illumina, 150 bp reads; Microsynth). NGS blast files were further processed by using tools on the Galaxy collaborative platform (https://galaxyproject.org). In brief, sequences were extracted according to their barcodes (“barcode splitter”), barcodes removed (“cutadapt”), and genome alignment (“Bowtie2”). The results were visualized via IGV (integrative genomic viewer, ver. 2.18.4)(75). The homology of genes to *SPD_0642* was analyzed via OrthoDB (version 12.1; https://sparql.orthodb.org/)(76).

### Statistics

GraphPad Prism 10 software (GraphPad Software, San Diego, CA, USA) was used for statistical analysis and graphics. A two-tailed p value of ≤0.05 was considered significant for all tests (α = 0.05). When two experimental groups were compared, the Mann‒Whitney U test was used. For comparisons of more than two groups, Kruskal–Wallis with Dunn’s multiple comparisons was used. For correlation analysis, both Pearson’s r (considered highly correlated when r>0.7) and p (considered significant at <0.05) were used. For the animal experiments, the sample size was determined via G-power software (version 3.1.9.6; Franz Faul, University of Kiel, Germany). For survival curve analysis, log-rank (Mantel-Cox) test and Gehan-Breslow-Wilcoxon test were used. No data were excluded, and the animals were randomized such that the male/female ratio in each group was close to equal.

## Data availability

Accession numbers for the presented sequences: D39 (accession number GCF_023635025.1), 106.66 (accession number PRJNA554545), 50943, 51060, 807.13, 51113, 51114, 48294, and 50793 (available upon publication). All experimental data will be made publicly available upon publication.

## Acknowledgments

This work was supported by grant No. 170844 from the Swiss–South African Joint Research Programme funded by the Swiss National Science Foundation and the National Research Foundation of South Africa to LJH and AvG, grant No. 160136 by the Swiss National Science Foundation to AII and Novartis Foundation Grant No. 20A010 to AII. The funding body had no role in the design of the study or collection, analysis or interpretation of the data or writing of the manuscript.

We are grateful to Susanne Aebi and Lalaina Holivololona for excellent technical assistance. We thank Prof. Alban Ramette and the NGS team from the Institute for Infectious Diseases (IFIK), Bern, for their technical help with some of the NGS experiments.

## Author contributions statement

Conceived the study and designed the experiments: LJH, AII, AVG, AM, SH. Performed the experiments: AM, SH, IT, AII, DB, NT, JL, CL. Materials were provided by FR and TvO. Analyzed the data: AM, SH, AII, CL and FR. Wrote the manuscript: AM, SH, LJH, and AII. All the authors discussed and approved the results presented in this manuscript.

## Ethics statement

Sample collection was performed with approval from the Cantonal Ethics Commission (Bern, Switzerland) and the Human Research Ethics Committee (Medical) of the University of Witwatersrand (Johannesburg, South Africa). The relevant university and provincial ethics committees approved the GERMS-SA surveillance study (clearance nos. M140159, M081117, M021042, and M180101). Animal experiments were performed in accordance with the Bern cantonal and Swiss national legislation, permits were obtained under No. BE103/2020 and BE91/2025.

## References

1. Schmidt H, Heimann B, Djukic M, Mazurek C, Fels C, Wallesch CW, Nau R. 2006. Neuropsychological sequelae of bacterial and viral meningitis. Brain 129:333–345.

2. Saez-Llorens X, McCracken Jr GH. 2003. Bacterial meningitis in children. Lancet. 361:2139–2148.

3. Mwenda JM, Soda E, Weldegebriel G, Katsande R, Biey JNM, Traore T, De Gouveia L, Du Plessis M, Von Gottberg A, Antonio M, Kwambana-Adams B, Worwui A, Gierke R, Schwartz S, Van Beneden C, Cohen A, Serhan F, Lessa FC. 2019. Pediatric Bacterial Meningitis Surveillance in the World Health Organization African Region Using the Invasive Bacterial Vaccine-Preventable Disease Surveillance Network, 2011-2016. Clin Infect Dis 69:S49–S57.

4. van Aalst M, Lötsch F, Spijker R, van der Meer JTM, Langendam MW, Goorhuis A, Grobusch MP, de Bree GJ. 2018. Incidence of invasive pneumococcal disease in immunocompromised patients: A systematic review and meta-analysis. Travel Med Infect Dis 24:89–100.

5. Tenforde MW, Gertz AM, Lawrence DS, Wills NK, Guthrie BL, Farquhar C, Jarvis JN. 2020. Mortality from HIV-associated meningitis in sub-Saharan Africa: a systematic review and meta-analysis. J Int AIDS Soc 23.

6. Kleynhans J, Cohen C, McMorrow M, Tempia S, Crowther-Gibson P, Quan V, de Gouveia L, von Gottberg A. 2019. Can pneumococcal meningitis surveillance be used to assess the impact of pneumococcal conjugate vaccine on total invasive pneumococcal disease? A case-study from South Africa, 2005–2016. Vaccine 37:5724–5730.

7. Ganaie F, Saad JS, McGee L, van Tonder AJ, Bentley SD, Lo SW, Gladstone RA, Turner P, Keenan JD, Breiman RF, Nahm MH. 2020. A New Pneumococcal Capsule Type, 10D, is the 100th Serotype and Has a Large cps Fragment from an Oral Streptococcus. mBio 11.

8. Martens P, Worm SW, Lundgren B, Konradsen HB, Benfield T. 2004. Serotype-specific mortality from invasive Streptococcus pneumoniae disease revisited. BMC Infect Dis 4:21.

9. Rückinger S, Von Kries R, Siedler A, Van Der Linden M. 2009. Association of serotype of Streptococcus pneumoniae with risk of severe and fatal outcome. Pediatr Infect Dis J 28:118–122.

10. Hathaway LJ, Brugger SD, Morand B, Bangert M, Rotzetter JU, Hauser C, Graber WA, Gore S, Kadioglu A, Mühlemann K. 2012. Capsule Type of Streptococcus pneumoniae Determines Growth Phenotype 8:e1002574.

11. Müller A, Salmen A, Aebi S, De Gouveia L, Von Gottberg A, Hathaway LJ. 2020. Pneumococcal serotype determines growth and capsule size in human cerebrospinal fluid. BMC Microbiol 20:1–9.

12. Hathaway LJ, Grandgirard D, Valente LG, Täuber MG, Leib SL. 2016. Streptococcus pneumoniae capsule determines disease severity in experimental pneumococcal meningitis. Open Biol 6.

13. Li J, Zhang J-R. 2019. Phase Variation of Streptococcus pneumoniae. Microbiol Spectr 7.

14. Li J, Li JW, Feng Z, Wang J, An H, Liu Y, Wang Y, Wang K, Zhang X, Miao Z, Liang W, Sebra R, Wang G, Wang WC, Zhang JR. 2016. Epigenetic Switch Driven by DNA Inversions Dictates Phase Variation in Streptococcus pneumoniae. PLoS Pathog 12.

15. Talbot UM, Paton AW, Paton JC. 1996. Uptake of Streptococcus pneumoniae by respiratory epithelial cells. Infect Immun1996/09/01. 64:3772–3777.

16. Weiser JN, Austrian R, Sreenivasan PK, Masure HR. 1994. Phase variation in pneumococcal opacity: relationship between colonial morphology and nasopharyngeal colonization. Infect Immun 62:2582–2589.

17. Li JW, Li J, Wang J, Li C, Zhang JR. 2019. Molecular Mechanisms of hsdS Inversions in the cod Locus of Streptococcus pneumoniae. J Bacteriol 201.

18. Shainheit MG, Mulé M, Camilli A. 2014. The Core Promoter of the Capsule Operon of Streptococcus pneumoniae Is Necessary for Colonization and Invasive Disease. Infect Immun 82:694.

19. Glanville DG, Gazioglu O, Marra M, Tokars VL, Kushnir T, Habtom M, Croucher NJ, Nebenzahl YM, Mondragón A, Yesilkaya H, Ulijasz AT. 2023. Pneumococcal capsule expression is controlled through a conserved, distal cis-regulatory element during infection. PLoS Pathog 19:e1011035.

20. Hirst RAA, Gosai B, Rutman A, Guerin CJJ, Nicotera P, Andrew PWW, O’Callaghan C, O’Callaghan C. 2008. Streptococcus pneumoniae Deficient in Pneumolysin or Autolysin Has Reduced Virulence in Meningitis. J Infect Dis 197:744–751.

21. Iliev AI, Djannatian JR, Nau R, Mitchell TJ, Wouters FS. 2007. Cholesterol-dependent actin remodeling via RhoA and Rac1 activation by the Streptococcus pneumoniae toxin pneumolysin. Proc Natl Acad Sci U S A. 104:2897–2902.

22. Hupp S, Heimeroth V, Wippel C, Fortsch C, Ma J, Mitchell TJ, Iliev AI. 2012. Astrocytic tissue remodeling by the meningitis neurotoxin pneumolysin facilitates pathogen tissue penetration and produces interstitial brain edema. Glia. 60:137–146.

23. Hupp S, Förtsch C, Graber F, Mitchell TJ, Iliev AI. 2022. Pneumolysin boosts the neuroinflammatory response to Streptococcus pneumoniae through enhanced endocytosis. Nature Communications 2022 13:1 13:1–18.

24. Laborada G, Rego M, Jain A, Guliano M, Stavola J, Ballabh P, Krauss AN, Auld PAM, Nesin M. 2003. Diagnostic value of cytokines and C-reactive protein in the first 24 hours of neonatal sepsis. Am J Perinatol 20:491–501.

25. Mook-Kanamori B, Geldhoff M, Troost D, van der Poll T, van de Beek D. 2012. Characterization of a pneumococcal meningitis mouse model. BMC Infect Dis 12:71.

26. Müller A, Kleynhans J, de Gouveia L, Meiring S, Cohen C, Hathaway LJ, von Gottberg A. 2022. Streptococcus pneumoniae Serotypes Associated with Death, South Africa, 2012-2018. Emerg Infect Dis 28:166–179.

27. Grabenstein JD, Musey LK. 2014. Differences in serious clinical outcomes of infection caused by specific pneumococcal serotypes among adults. Vaccine 32:2399–2405.

28. Olarte L, Kaplan SL, Barson WJ, Romero JR, Lin PL, Tan TQ, Hoffman JA, Bradley JS, Givner LB, Mason EO, Hultén KG. 2017. Emergence of multidrug-resistant pneumococcal serotype 35B among children in the United States. J Clin Microbiol 55:724–734.

29. King LM, Andrejko KL, Kobayashi M, Xing W, Cohen AL, Self WH, Resser JJ, Whitney CG, Baughman A, Kio M, Grijalva CG, Traenkner J, Rouphael N, Lewnard JA. 2024. PNEUMOCOCCAL SEROTYPE DISTRIBUTION AND COVERAGE OF EXISTING AND PIPELINE PNEUMOCOCCAL VACCINES. medRxiv 2024.12.12.24318944.

30. Seeberger PH, Pereira CL, Govindan S. 2017. Total synthesis of a Streptococcus pneumoniae serotype 12F CPS repeating unit hexasaccharide. Beilstein Journal of Organic Chemistry 13:164.

31. Weiser JN, Bae D, Epino H, Gordon SB, Kapoor M, Zenewicz LA, Shchepetov M. 2001. Changes in Availability of Oxygen Accentuate Differences in Capsular Polysaccharide Expression by Phenotypic Variants and Clinical Isolates of Streptococcus pneumoniae. Infect Immun 69:5430.

32. Schaffner TO, Hinds J, Gould KA, Wüthrich D, Bruggmann R, Küffer M, Mühlemann K, Hilty M, Hathaway LJ. 2014. A point mutation in cpsE renders Streptococcus pneumoniae nonencapsulated and enhances its growth, adherence and competence. BMC Microbiol 14:210.

33. Carvalho SM, Kloosterman TG, Manzoor I, Caldas J, Vinga S, Martinussen J, Saraiva LM, Kuipers OP, Neves AR. 2018. Interplay between capsule expression and uracil metabolism in Streptococcus pneumoniae D39. Front Microbiol 9:319383.

34. Geisinger E, Isberg RR. 2015. Antibiotic Modulation of Capsular Exopolysaccharide and Virulence in Acinetobacter baumannii. PLoS Pathog 11:e1004691.

35. Sakatani H, Kono M, Sugita G, Nanushaj D, Hijiya M, Iyo T, Shiga T, Murakami D, Kaku N, Yanagihara K, Nahm MH, Hotomi M. 2022. Investigation on the virulence of non-encapsulated Streptococcus pneumoniae using liquid agar pneumonia model. Journal of Infection and Chemotherapy 28:1452–1458.

36. Keller LE, Robinson DA, McDaniel LS. 2016. Nonencapsulated Streptococcus pneumoniae: Emergence and Pathogenesis. mBio 7:e01792–15.

37. Bradshaw JL, McDaniel LS. 2019. Selective pressure: Rise of the nonencapsulated pneumococcus. PLoS Pathog 15.

38. Hu L, Shi Y, Xu Q, Zhang L, He J, Jiang Y, Liu L, Leptihn S, Yu Y, Hua X, Zhou Z. 2020. Capsule Thickness, Not Biofilm Formation, Gives Rise to Mucoid Acinetobacter baumannii Phenotypes That are More Prevalent in Long-Term Infections: A Study of Clinical Isolates from a Hospital in China. Infect Drug Resist 13:99–109.

39. Rakovitsky N, Lellouche J, Ben David D, Frenk S, Elmalih P, Weber G, Kon H, Schwartz D, Wolfhart L, Temkin E, Carmeli Y. 2021. Increased Capsule Thickness and Hypermotility Are Traits of Carbapenem-Resistant Acinetobacter baumannii ST3 Strains Causing Fulminant Infection. Open Forum Infect Dis 8.

40. Zafar MA, Hamaguchi S, Zangari T, Cammer M, Weiser JN. 2017. Capsule type and amount affect shedding and transmission of streptococcus pneumoniae. mBio 8.

41. Nelson AL, Roche AM, Gould JM, Chim K, Ratner AJ, Weiser JN. 2007. Capsule enhances pneumococcal colonization by limiting mucus-mediated clearance. Infect Immun 75:83–90.

42. Zhu J, Abruzzo AR, Wu C, Bee GCW, Pironti A, Putzel G, Aggarwal SD, Eichner H, Weiser JN. 2023. Effects of Capsular Polysaccharide amount on Pneumococcal-Host interactions. PLoS Pathog 19:e1011509.

43. Hammerschmidt S, Wolff S, Hocke A, Rosseau S, Müller E, Rohde M, Muller E, Rohde M. 2005. Illustration of Pneumococcal Polysaccharide Capsule during Adherence and Invasion of Epithelial Cells. Infect Immun 73.

44. Tanimoto H, Shigemura K, Osawa K, Kado M, Onishi R, Fang S Bin, Sung SY, Miyara T, Fujisawa M. 2023. Comparative genetic analysis of the antimicrobial susceptibilities and virulence of hypermucoviscous and non-hypermucoviscous ESBL-producing Klebsiella pneumoniae in Japan. Journal of Microbiology, Immunology and Infection 56:93–103.

45. Gendrin C, Merillat S, Vornhagen J, Coleman M, Armistead B, Ngo L, Aggarwal A, Quach P, Berrigan J, Rajagopal L. 2018. Diminished Capsule Exacerbates Virulence, Blood–Brain Barrier Penetration, Intracellular Persistence, and Antibiotic Evasion of Hyperhemolytic Group B Streptococci. J Infect Dis 217:1128–1138.

46. Quessy S, Dubreuil JD, Jacques M, Malouin F, Higgins R. 1994. Increase of capsular material thickness following in vivo growth of virulent Streptococcus suis serotype 2 strains. FEMS Microbiol Lett 115:19–26.

47. Moorthy AN, Rai P, Jiao H, Wang S, Tan KB, Qin L, Watanabe H, Zhang Y, Narasaraju T, Chow VTK, Moorthy AN, Rai P, Jiao H, Wang S, Tan KB, Qin L, Watanabe H, Zhang Y, Narasaraju T, Chow VTK. 2016. Capsules of virulent pneumococcal serotypes enhance formation of neutrophil extracellular traps during in vivo pathogenesis of pneumonia. Oncotarget 7:19327–19340.

48. Henke MT, Brown EM, Cassilly CD, Vlamakis H, Xavier RJ, Clardy J. 2021. Capsular polysaccharide correlates with immune response to the human gut microbe Ruminococcus gnavus. Proc Natl Acad Sci U S A 118:e2007595118.

49. Loh E, Kugelberg E, Tracy A, Zhang Q, Gollan B, Ewles H, Chalmers R, Pelicic V, Tang CM. 2013. Temperature triggers immune evasion by Neisseria meningitidis. Nature 502:10.1038/nature12616.

50. Buffet A, Rocha EPC, Rendueles O. 2021. Nutrient conditions are primary drivers of bacterial capsule maintenance in Klebsiella. Proceedings of the Royal Society B 288.

51. Manso AS, Chai MH, Atack JM, Furi L, De Ste Croix M, Haigh R, Trappetti C, Ogunniyi AD, Shewell LK, Boitano M, Clark TA, Korlach J, Blades M, Mirkes E, Gorban AN, Paton JC, Jennings MP, Oggioni MR. 2014. A random six-phase switch regulates pneumococcal virulence via global epigenetic changes. Nat Commun 5.

52. Troxler LJ, Werren JP, Schaffner TO, Mostacci N, Vermathen P, Vermathen M, Wüthrich D, Simillion C, Brugger SD, Bruggmann R, Hathaway LJ, Furrer J, Hilty M. 2019. Carbon source regulates polysaccharide capsule biosynthesis in Streptococcus pneumoniae. Journal of Biological Chemistry 294:17224–17238.

53. Echenique J, Kadioglu A, Romao S, Andrew PW, Trombe MC. 2004. Protein serine/threonine kinase StkP positively controls virulence and competence in Streptococcus pneumoniae. Infect Immun 72:2434–2437.

54. Morona JK, Paton JC, Miller DC, Morona R. 2000. Tyrosine phosphorylation of CpsD negatively regulates capsular polysaccharide biosynthesis in streptococcus pneumoniae. Mol Microbiol 35:1431–1442.

55. O’Meara TR, Andrew Alspaugh J. 2012. The Cryptococcus neoformans capsule: A sword and a shield. Clin Microbiol Rev 25:387–408.

56. Lai Y-C, Peng H-L, Chang H-Y. 2003. RmpA2, an Activator of Capsule Biosynthesis in Klebsiella pneumoniae CG43, Regulates K2 cps Gene Expression at the Transcriptional Level. J Bacteriol 185:788–800.

57. Campos MA, Vargas MA, Regueiro V, Llompart CM, Albertí S, Bengoechea JA. 2004. Capsule Polysaccharide Mediates Bacterial Resistance to Antimicrobial Peptides. Infect Immun 72:7107.

58. Geisinger E, Isberg RR. 2015. Antibiotic Modulation of Capsular Exopolysaccharide and Virulence in Acinetobacter baumannii. PLoS Pathog 11:e1004691.

59. Geisinger E, Isberg RR. 2015. Antibiotic Modulation of Capsular Exopolysaccharide and Virulence in Acinetobacter baumannii. PLoS Pathog 11:e1004691.

60. Zheng Y, Zhang X, Wang X, Wang L, Zhang J, Yin Y. 2017. ComE, an essential response regulator, negatively regulates the expression of the capsular polysaccharide locus and attenuates the bacterial virulence in Streptococcus pneumoniae. Front Microbiol 8:234866.

61. Zheng Y-D, Pan Y, He K, Li N, Yang D, Du G-F, Ge R, He Q-Y, Sun X. 2020. SPD_1495 Contributes to Capsular Polysaccharide Synthesis and Virulence in Streptococcus pneumoniae . mSystems 5.

62. Junges R, Salvadori G, Shekhar S, Åmdal HA, Periselneris JN, Chen T, Brown JS, Petersen FC. 2017. A Quorum-Sensing System That Regulates Streptococcus pneumoniae Biofilm Formation and Surface Polysaccharide Production. mSphere 2.

63. Hathaway LJ, Bättig P, Reber S, Rotzetter JU, Aebi S, Hauser C, Heller M, Kadioglu A, Mühlemann K. 2014. Streptococcus pneumoniae detects and responds to foreign bacterial peptide fragments in its environment. Open Biol 4.

64. Nasher F, Heller M, Hathaway LJ. 2018. Streptococcus pneumoniae Proteins AmiA, AliA, and AliB Bind Peptides Found in Ribosomal Proteins of Other Bacterial Species. Front Microbiol 8.

65. Nasher F, Aguilar F, Aebi S, Hermans PWM, Heller M, Hathaway LJ. 2018. Peptide Ligands of AmiA, AliA, and AliB Proteins Determine Pneumococcal Phenotype. Front Microbiol 9.

66. Wu K, Xu H, Zheng Y, Wang L, Zhang X, Yin Y. 2016. CpsR, a GntR family regulator, transcriptionally regulates capsular polysaccharide biosynthesis and governs bacterial virulence in Streptococcus pneumoniae. Sci Rep 6.

67. Zhang J, Ye W, Wu K, Xiao S, Zheng Y, Shu Z, Yin Y, Zhang X. 2021. Inactivation of Transcriptional Regulator FabT Influences Colony Phase Variation of Streptococcus pneumoniae. mBio 12.

68. Ferrada E, Superti-Furga G. 2022. A structure and evolutionary-based classification of solute carriers. iScience 25:105096.

69. Frommelt F, Ladurner R, Goldmann U, Wolf G, Ingles-Prieto A, Lineiro-Retes E, Gelová Z, Hopp AK, Christodoulaki E, Teoh ST, Leippe P, Santini BL, Rebsamen M, Lindinger S, Serrano I, Onstein S, Klimek C, Barbosa B, Pantielieieva A, Dvorak V, Hannich TJ, Schoenbett J, Sansig G, Mocking TAM, Ooms JF, IJzerman AP, Heitman LH, Sykacek P, Reinhardt J, Müller AC, Wiedmer T, Superti-Furga G. 2025. The solute carrier superfamily interactome. Molecular Systems Biology 21:632–675.

70. Mühlemann K, Matter HC, Täuber MG, Bodmer T. 2003. Nationwide surveillance of nasopharyngeal Streptococcus pneumoniae isolates from children with respiratory infection, Switzerland, 1998-1999. J Infect Dis 187:589–596.

71. van Opijnen T, Lazinski DW, Camilli A. 2014. Genome-Wide Fitness and Genetic Interactions Determined by Tn-seq, a High-Throughput Massively Parallel Sequencing Method for Microorganisms. Curr Protoc Mol Biol 106:7.16.1-7.16.24.

72. Gates MA, Thorkildson P, Kozel TR. 2004. Molecular architecture of the Cryptococcus neoformans capsule. Mol Microbiol 52:13–24.

73. Chiavolini D, Tripodi S, Parigi R, Oggioni MR, Blasi E, Cintorino M, Pozzi G, Ricci S. 2004. Method for inducing experimental pneumococcal meningitis in outbred mice, p. 36. In BMC Microbiol. BioMed Central, England.

74. Li J, Wang J, Jiao F, Zhang J-R. 2017. Observation of Pneumococcal Phase Variation in Colony Morphology. Bio Protoc 7:e2434.

75. Robinson JT, Thorvaldsdottir H, Turner D, Mesirov JP. 2023. igv.js: an embeddable JavaScript implementation of the Integrative Genomics Viewer (IGV). Bioinformatics 39.

76. Tegenfeldt F, Kuznetsov D, Manni M, Berkeley M, Zdobnov EM, Kriventseva E V. 2025. OrthoDB and BUSCO update: annotation of orthologs with wider sampling of genomes. Nucleic Acids Res 53:D516–D522.

